# Early seed priming with closely related *Bacillus* strains induces divergent physiological and defense responses in melon

**DOI:** 10.1101/2025.07.16.665077

**Authors:** Luisa Carrégalo-Ríos, Carlos Molina-Santiago, María V. Berlanga-Clavero, Daniel Petras, Jesús Hierrezuelo, Mónica Pineda, Juan M. Alba, Antonio de Vicente, Matilde Barón-Ayala, Pieter C. Dorrestein, Diego Romero

## Abstract

Early microbial seed priming is a promising strategy to improve crop resilience, yet it remains unclear whether plants can discriminate among closely related beneficial strains and integrate dose-dependent microbial cues. We primed melon (*Cucumis melo*) seeds with two phylogenetically similar *Bacillus* strains (*B. subtilis* NCIB3610 and *B. velezensis* FZB42) and combined transcriptomic, metabolomic and physiological analyses across development. Despite comparable colonization, the strains provoked contrasting host programs and distinct dose responses. *B. subtilis* promoted radicle elongation, chloroplastic starch storage and drought tolerance irrespective of inoculum level, together with L-tryptophan and palatinose accumulation. By contrast, *B. velezensis* displayed a clear dose effect: low inoculum sustained normal radicle growth, whereas high inoculum transiently repressed it, coinciding with retrotransposon activation, suppression of AOS and proteasome genes, and enrichment of flavonoids and glutathione in leaves. Chemical assays showed that radicle inhibition requires the combination of surfactin, produced by both strains, with bacillomycin D, an iturin-type lipopeptide unique to FZB42; neither molecule alone reproduced the effect. This synergy links the strain-specific lipopeptide repertoire to the dose-dependent growth response. Although their early trajectories diverged, both primings converged on improved above-ground performance. 3610-primed plants restricted *Botrytis cinerea* via caffeic- and rosmarinic-acid accumulation, whereas FZB42-primed plants curtailed jasmonate-sensitive *Tetranychus urticae* mites through early JA-pathway activation. Our results demonstrate that melon perceives inoculum dose and microbial identity, translating them into distinct metabolic and defense programs that converge on stress resilience. These mechanistic insights-linking lipopeptide fingerprints, sentinel metabolites and defense transcripts-provide a framework for precision seed treatments in horticultural crops.

## Introduction

Plants have coevolved with diverse microbial partners, forming interactions that range from mutualistic to pathogenic^1,2^. The ability of plants to perceive and respond appropriately to microbial cues is central to maintaining this ecological balance, shaped by evolutionary selection of those interactions that promote survival and fitness of the plant holobiont^3^. While much attention has focused on defense responses to pathogens, less is known about how plants differentially interpret beneficial microbes, particularly those that are closely related yet exert a distinct function. Understanding how plants discriminate among beneficial microorganisms is key to unraveling the ecological and evolutionary dynamics of plant–microbiome interactions. Microbial associations initiated at the seed stage are among the earliest interactions in the plant life cycle^4,5^ and can influence development and stress responses across tissues and life stages^6^. These early interactions offer a unique opportunity to explore how plants integrate microbial information to shape their adaptive trajectory throughout their development.

*Bacillus* species are widely used as plant growth–promoting rhizobacteria^7,8^, however, most studies treat these microbes as functionally equivalent, focusing on their general effects on yield or pathogen suppression^9^. Much of the literature focuses on a single strain and its ability to promote growth or enhance stress tolerance^10^, overlooking the underlying recognition mechanisms that might lead to distinct, yet beneficial responses that could converge on a similar phenotype. From the plant’s perspective, mutualism is not a binary state, but a flexible interaction shaped by functional compatibility. Different beneficial microbes, even those that are phylogenetically close, may engage the host in distinct ways^11,12^. The extent to which a plant can modulate its metabolic and defense programs in response to strain-specific interactions remains largely unexplored.

In this study, we use early seed priming with two closely related *Bacillus* strains, *B. subtilis* NCIB3610 and *B. velezensis* FZB42, as a tool to probe how the plant host modulates its adaptive programming in a mutualistic context. Despite similar colonization efficiency, the two strains trigger distinct developmental, metabolic, and defensive trajectories in *Cucumis melo*. By integrating transcriptomic, metabolomic, anatomical and physiological data, we show that the plant does not merely respond to microbial presence, but actively differentiates between beneficial partners, demonstrating that mutualism itself can take multiple functional forms, shaped by microbial identity.

## Results

### Dose-dependent root developmental responses to *Bacillus* seed treatments

Melon seeds treated with *Bacillus velezensis* FZB42 exhibited smaller radicles compared to those treated with either *Bacillus subtilis* NCIB3610 or distilled water (referred to as “mock plants” hereafter) after 5 days of growth (Fig. 1A,B). Both bacterial strains were able to colonize the seed and persisted during radicle development, reaching 100% sporulation within 48 hours (Fig. S1A). *B. subtilis and B. velezensis* were also recovered from adult plant roots, with sporulation rates of 31.2% and 57.3%, respectively (Fig. S1B), highlighting their successful establishment as root colonizers. To understand the observed differences in radicle growth, various bacterial inoculum concentrations were tested. A lower inoculum concentration of *B. velezensis* (10^7^ colony-forming unit (CFU)/ml instead of the initial 10^8^ CFU/ml) restored radicle growth to mock plant phenotype, while *B. subtilis* consistently showed a trend towards growth promotion at both doses tested (Fig. 1C). The radicle growth repression induced by *B. velezensis* at higher inoculum concentration was also observed in *Arabidopsis thaliana*, a species phylogenetically distant from melon (Fig. S1C). Importantly, the early repression of radicle growth did not translate into detrimental consequences during later developmental stages, as all plants reached the same developmental stage by 35 days (Fig. 1A,B).

**Figure 1.**
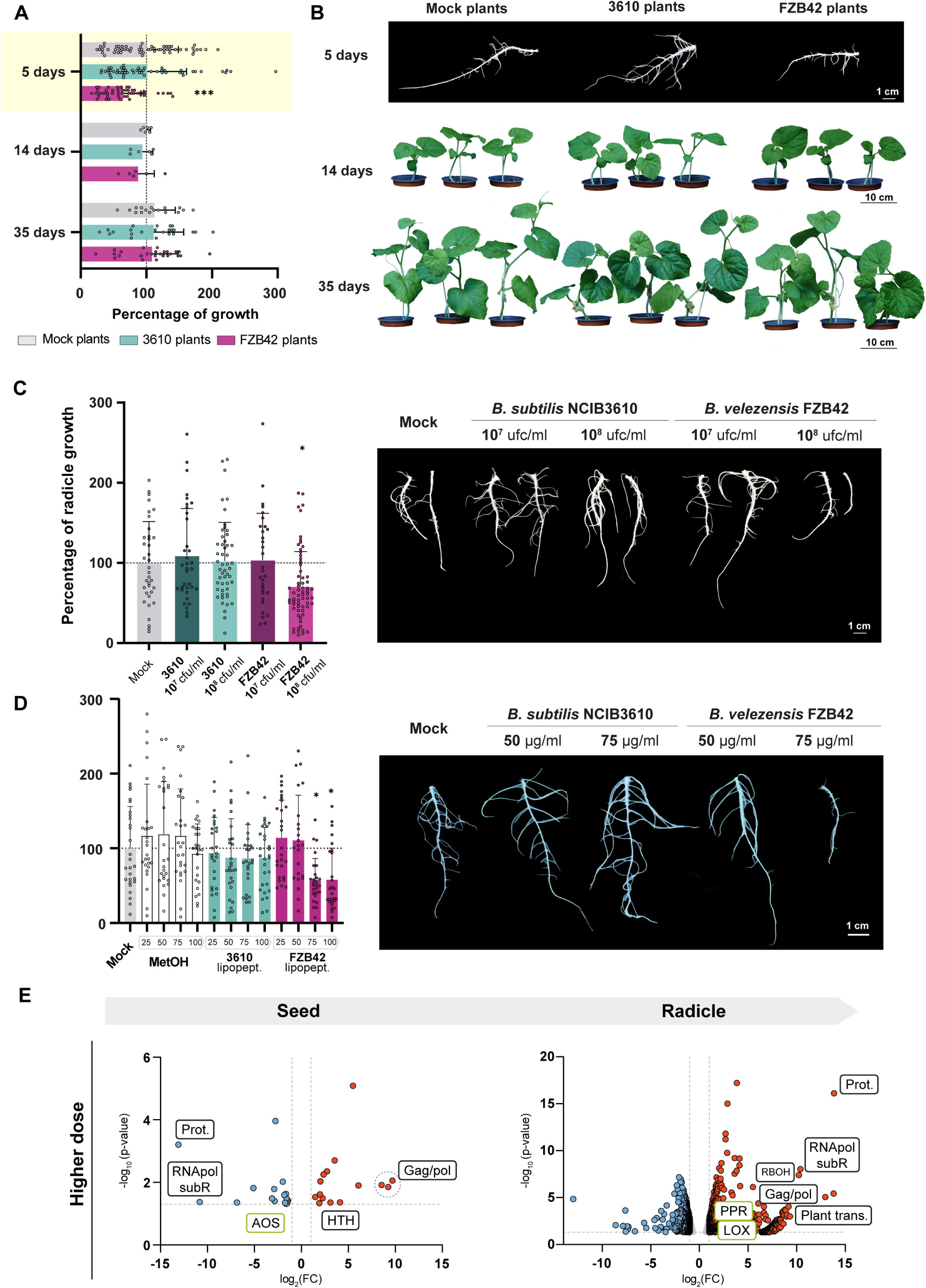
Early seed priming with *B. velezensis* delays development despite exhibiting similar ecological traits to *B. subtilis* on melon seeds. **a** Growth progression (%) of plants grown from seeds treated with *B. subtilis* NCIB3610 (blue) or *B. velezensis* FZB42 (purple), relative to mock-treated controls (gray, distilled water). Growth was normalized to the average radicle area at 5 days post treatment (dpt) (highlighted, yellow background) or average fresh weight at 14 and 35 dpt (white background) of mock plants (dashed line = 100%). Mean ± SD. Statistical significance was assessed using one-way ANOVA with Dunnett’s post-hoc test (***P = 0.0002 vs. mock). **b** Representative images of radicles (5 dpt) and whole plants (14 and 35 dpt) derived from mock- and *Bacillus*-treated seeds. **c** Left: Radicle development (%) 5 days after seed inoculation with *B. subtilis* or *B. velezensis* at low (10^7^ CFU/ml) or high (10^8^ CFU/ml) bacterial densities. Data are normalized to mock controls (dashed line = 100%). Mean ± SD. Statistical analysis as above (*P < 0.01). Right: Representative radicles from each treatment. Scale bar = 1 cm. **d** Left: Radicle development (%) 5 dpt with crude lipopeptide extracts from *B. subtilis* or *B. velezensis* (25–100 µg/ml). Methanol control included. Data normalized to mock controls (dashed line = 100%). Mean ± SD. One-way ANOVA with Dunnett’s test (*P < 0.01). Right: Representative radicles per condition. Scale bar = 1 cm. **e** Volcano plot showing differentially expressed genes (DEGs) in seeds treated with high-dose *B. velezensis* and their corresponding radicles. P-values calculated using Fisher’s method across edgeR and DESeq2 results. Thresholds for significance (horizontal = P, vertical = fold change) are shown as dashed lines. Green labels indicate DEGs shared with low inoculum dataset. Selected genes are annotated for retrotransposon activity, oxidative stress, and plant defense (full data in Table S1).

Previous report indicated that fengycin contributes to oxidative stress and tissue damage when added to melon seeds, leading to overgrowth of adult plants^13^. In addition to fengycin, *B. velezensis* FZB42, but not *B. subtilis* NCIB3610, produces bacillomycin D, an iturin-like lipopeptide (Fig. S1D)^14^, which led to speculate that exposure to this compound could have a negative effect on root growth. Supporting this, when crude lipopeptide extracts from bacterial cultures were applied to melon seeds, only extracts from *B. velezensis* at 75 µg/ml and 100 µg/ml induced significant radicle growth inhibition (Fig. 1D). Lower concentrations (25 and 50 µg/ml) did not reproduce the effect. Interestingly, treatment with purified single lipopeptides failed to inhibit radicle growth. Bacillomycin D (20 µM) even promoted growth, both alone and co-inoculated with fengycin, whereas surfactin (20 µM) showed a tendency toward repression. Notably, the growth-promoting effects of bacillomycin D and fengycin were lost when surfactin was included in the treatment (Fig. S1E). These findings suggest a synergistic mechanism among lipopeptides or structural variants, contributing to growth repression.

We previously reported that initial radicle growth does not necessarily predict adult plant development or resistance to biotic stress. Nevertheless, *B. subtilis* triggered changes in the dynamics of triacylglycerides after differential targeting of seed oil bodies and lipid catabolism optimization, as shown by transcriptomic analysis^13^. Since all groups reached similar developmental stages over time, we hypothesized that seeds treated with a higher inoculum of *B. velezensis* ("FZB42 plants") undergo adaptive developmental reprogramming, favoring long-term benefits over early growth. Transcriptomic analysis performed immediately after seed treatment with high inoculum of *B. velezensis* revealed strong upregulation of endogenous retrovirus and retrotransposon-related genes, particularly those encoding Gag/Pol proteins (Fig. 1E), alongside repression of proteasome components (Table S1), possibly stabilizing retrotransposon activity. Transposable elements are known to influence genome architecture and gene regulation in response to environmental stimuli^15^. Consistent with this, radicle transcriptomes from high inoculum treatments showed globally increased gene expression levels (Fig. 1E, Fig. S2A). A particularly notable result was the downregulation of allene oxide synthase (AOS), a key enzyme in jasmonic acid (JA) biosynthesis^16^, with a stronger effect at high dose (log_2_FC = -2.9) compared to the lower dose (log_2_FC = -1.7; Table S1). In the context of the JAZ repressor model, where low JA-Ile levels maintain a primed defense state^17^, this downregulation may lower the activation threshold for future defense responses.

Taken together, these findings suggest a potential epigenetic memory, in which early activation of transposable elements and gene expression reprogramming may prepare the plant for enhanced resilience against future stressors. The observed oxidative stress and transcriptional activity in high-dose radicles (Fig. 1E; Fig. S2B,C) support this model.

### Aboveground physiological and anatomical adaptations following *Bacillus* priming

Given the transcriptomic and phenotypic differences observed during seed and radicle stages, we analyzed metabolic profiles in adult leaves. Metabolomic analysis across leaf sections from mock plants revealed age-dependent variations, as well as metabolite accumulation influenced by seed treatment (Fig. 2A). The most pronounced changes were observed in older leaves of FZB42 plants (Fig. S3) with between 300 and 500 differentially accumulated metabolites in the second and third leaves. Younger leaves showed fewer than 100 accumulated metabolites. *B. subtilis*-treated plants (“3610 plants”) showed a similar gradient, with older leaves accumulating more metabolites than younger ones. Chemical classification of the top 50 metabolites differentially accumulated by treatment and leaf age revealed distinct signatures: FZB42 plants accumulated fatty acids, particularly in the oldest leaf, while 3610 plants exhibited increased levels in alkaloids, including tryptophan derivatives (Fig. 2B). Glycerolipids and glycerophospholipids were enriched in FZB42 plants, suggesting a shift in lipid metabolism (Fig. S4A).

**Figure 2.**
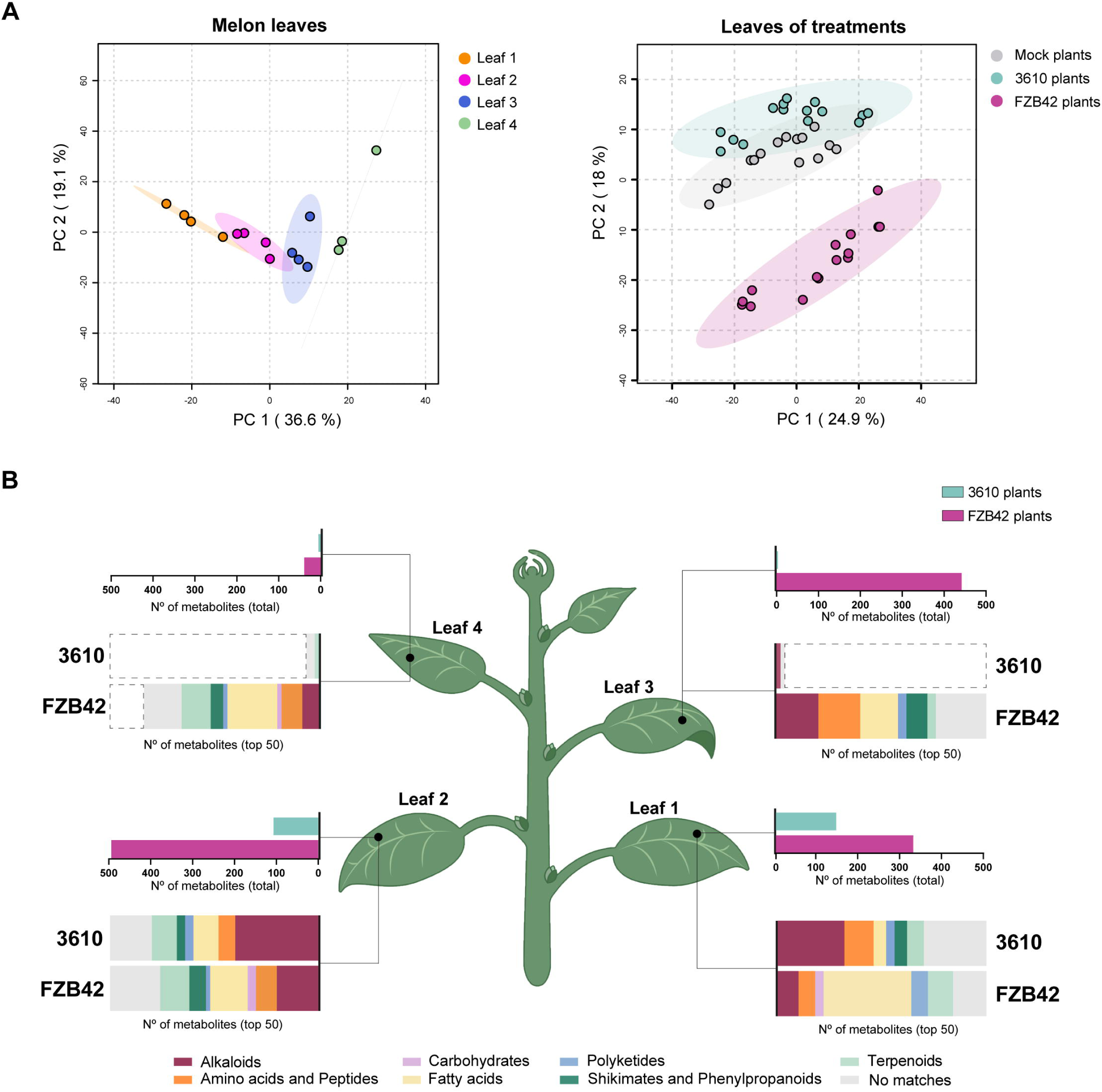
*Bacillus* strains trigger distinct patterns of metabolite accumulation in melon leaves. **a** Principal component analysis (PCA) of metabolomic profiles from melon leaves. Left: Clustering based on leaf developmental stage in mock-treated plants. Right: Clustering based on seed treatment (*B. subtilis* NCIB3610, *B. velezensis* FZB42, and mock). Percentage of variance explained by each principal component is indicated on the axes. **b** Metabolite accumulation summary based on volcano plot analysis comparing each *Bacillus* treatment to mock. Upper panels: number of significantly accumulated metabolites in 3610-treated (blue) and FZB42-treated (pink) plants. Lower panels: chemical classification (NPC#pathway) of the top 50 discriminant metabolites. Significance threshold: |log_2_(FC)| > 2 and FDR-adjusted P < 0.05.

All plant groups exhibited a shared healthy phenotype at the stage of adult plant, however, the transcriptomic (at seed and radicle level) and metabolic differences (leaves of adult plants) between treatments prompted us to investigate potential divergences in physiological and anatomical outcomes. Further refinement of feature-based molecular networking of the collected metabolome data identified three metabolites of interest: reduced glutathione (GSH), significantly accumulated in leaves of FZB42 plants; luteolin 7-glucoside, a flavonoid with potent antioxidant activity^18^, mainly accumulated in leaves of FZB42 plants but it was also present in 3610 plants; and protoporphyrin IX, accumulated in mock plants and not associated with either of the bacterial treatments (Fig. 3A). To investigate whether the reduced accumulation of a chlorophyll precursor in treated plants might compromise photosynthesis, a fundamental process for plant growth^19^, physiological assays were conducted. Photosynthetic performance of the second leaf was evaluated at 35 days after sowing (adult plants) by measuring the maximum quantum yield of photosystem II (PSII) (F_V_/F_M_) and effective quantum yield of PSII at steady-state (Φ_PSII_). In plants grown from *Bacillus*-treated seeds, imaging analysis revealed a slight reduction in photosynthetic efficiency at the leaf base near the petiole, particularly in terms of Φ_PSII_ compared to mock plants. However, this decrease was offset by increased photosynthetic activity in other leaf regions (Fig. 3B). Indeed, when averaged across the entire leaf surface, 3610 and FZB42 plants exhibited no significant differences in Φ_PSII_ or F_V_/F_M_ compared to mock plants, demonstrating that the overall photosynthetic yield of melon plants was not affected by seed treatment (Fig. 3C). Similarly, non-photochemical quenching at the steady-state (NPQ), representing energy not used in photosynthesis and dissipated as heat^20^, remained invariable between treatments (Fig. 3C). Photosynthesis is tightly regulated to prevent the accumulation of reactive intermediates that could induce oxidative stress^21^. Given the evidence of an elevated oxidative state in plants grown from *Bacillus*-treated seeds, marked by latent metabolic imprints in GSH and luteolin 7-glucoside accumulation, our results suggest an optimization of this fundamental metabolic process. Moreover, the greater water content within the leaves of 3610 and FZB42 plants compared to mock plants (Fig. 3D), further supports this hypothesis, in line with the well-established dependence of photosynthetic efficiency on leaf water status^22^. Finally, despite exhibiting less accumulation of Protoporphyrin IX, no differences in chlorophyll *a* and *b* content were observed in 3610 and FZB42 plants (Fig. 3E). In the light of these results, we hypothesize that 3610 and FZB42 plants might count with an ameliorated photosynthetic process where the same needs are met with fewer content of photosynthesis intermediates.

**Figure 3.**
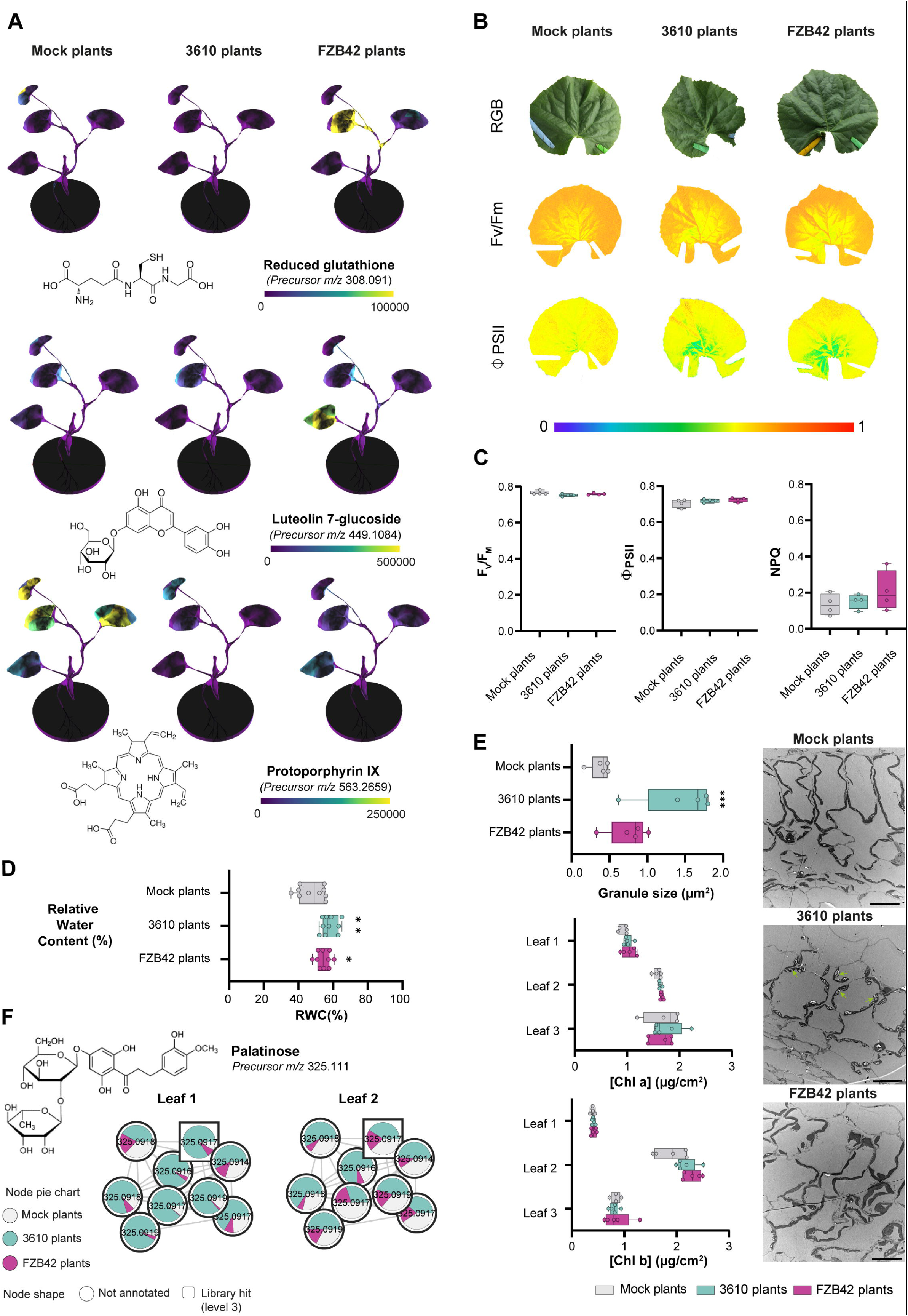
*Bacillus*-primed plants maintain photosynthetic capacity while showing adaptive traits including oxidative stress modulation and increased water retention. **a** 3D spatial distribution of selected metabolites in leaves of mock- and *Bacillus*-treated plants, identified by spectral match: reduced glutathione (top), luteolin 7-glucoside (middle), and protoporphyrin IX (bottom). **b** Representative images showing RGB, maximum quantum efficiency of PSII (F_V_/F_M_), and effective quantum yield of PSII (Φ_PSII_) in steady-state in leaves from each treatment group. **c** Quantitative values of F_V_/F_M_ (left), Φ_PSII_ (middle), and non-photochemical quenching (NPQ; right) from leaves of mock- and *Bacillus*-treated plants. Box plots show median (horizontal line) and range (whiskers). **d** Relative water content (RWC) of leaves from mock- and *Bacillus*-treated plants. Box and whisker plot. One-way ANOVA with Dunnett’s test (*P = 0.0313; **P = 0.0011). **e** Left: granule size (µm^2^) per plant group (top), chlorophyll *a* content (µg/cm^2^) by leaf age (middle), and chlorophyll *b* content (µg/cm^2^) by leaf age (bottom). Box and whisker plots based on five slices from three biological replicates. One-way ANOVA with Dunnett’s test (***P = 0.0005). Right: TEM images of thin leaf sections showing increased starch accumulation in chloroplasts of *B. subtilis*-primed plants. Scale bar = 20 µm. **f** Molecular family network of palatinose (isomaltulose) and related differentially abundant metabolites in first and second leaves. Pie charts represent average normalized abundance. Node shape indicates metabolite identification confidence(Sumner et al., 2007).

To visualize anatomical changes derived from seed treatment, second leaf sections of adult plants were analyzed by transmission electron microscopy (TEM) and scanning electron microscopy (SEM). Plants grown from seeds treated with *B. subtilis* exhibit increased starch accumulation inside chloroplasts, as shown by TEM analysis of thin leaf sections (Fig. 3E). Starch, a complex carbohydrate polymer commonly found in plant cells, accumulates during the day under light and breaks down at night to provide sugars for metabolism and growth^23^. Since all samples were collected at the same time of day, any differences observed between plant groups likely reflect variations in either the accumulation or degradation patterns of reserve carbohydrates. Based on non-targeted metabolomics data, palatinose (also known as isomaltulose; α-D- glucopyranosyl-1,2-D-fructofuranose)^24^ accumulates in older leaves of 3610 plants (Fig. 3F). This metabolite is a naturally occurring analog of sucrose, which has been reported to stimulate sucrose degradation and starch synthesis^25^.

Taken together, these findings suggest a potential regulatory interaction between palatinose accumulation and starch metabolism in 3610 plants, enabling enhanced starch synthesis in leaves. This mechanism likely facilitates the establishment of a carbohydrate reserve, ensuring both a readily available energy source for future metabolic demands and a source of soluble sugars that may function as osmoprotectants during environmental challenges^26^. Compared to mock and 3610 plants, FZB42 plants exhibited a higher stomatal density, but only on the adaxial surface of the second leaf at the adult plant stage (Fig. S5A,B). Additionally, FZB42 plants showed a tendency to maintain stomata open, suggesting an elevated leaf transpiration rate. Consistently, leaf surface temperature, inversely correlated with leaf transpiration and evaporation^27^, was significantly lower in FZB42 plants compared to the other treatments (Fig. S5C,D).

### Improved abiotic stress resilience linked to *Bacillus*-mediated seed priming

To explore potential landscapes where the forementioned physiological and metabolic differences may be significant, we assessed plant responses to various abiotic and biotic stressors. Given the increased water content in *Bacillus*-treated plants (Fig. 3D) and the potential role of starch in osmotic adjustment in 3610 plants (Fig. 3E), we investigated plant drought response. After 17 days without irrigation, 3610 and FZB42 plants appeared visibly healthier than mock plants, with 3610 plants exhibiting the highest relative leaf water content (RWC) of all group plants during stress conditions. Upon rewatering, 3610 and FZB42 plants recovered faster than mock plants; however, no significant differences in RWC were recorded at this stage (Fig. 4B). To further investigate the physiological basis of this response, we assessed photosynthetic performance during recovery by measuring F_V_/F_M_ and steady-state Φ_PSII_ across the entire second leaf surface (Fig. 4C,D). Compared to well-watered plants (Fig. 3B,C), all groups exhibited reduced photosynthetic activity, confirming that water deprivation severely impacted primary metabolism. However, 3610 and FZB42 plants showed significantly higher F_V_/F_M_ and steady-state Φ_PSII_ values than mock plants (Fig. 4D), suggesting that only the latter suffered irreversible damage to the chloroplast electron transport chain. Consistently, steady-state NPQ, a key photoprotective mechanism against photooxidation in the thylakoid membranes^28^, remained significantly elevated in *Bacillus*-treated plants in comparison to mock plants (Fig. 4D). Along with higher RWC and starch accumulation, L-tryptophan was significantly accumulated in both younger and older leaves of non-stressed 3610 plants, as well as in the third leaf of FZB42 plants (Fig. 4E). This amino acid may confer additional protection against desiccation, given that exogenous L-tryptophan treatment has been reported to alleviate hydric stress^29^.

**Figure 4.**
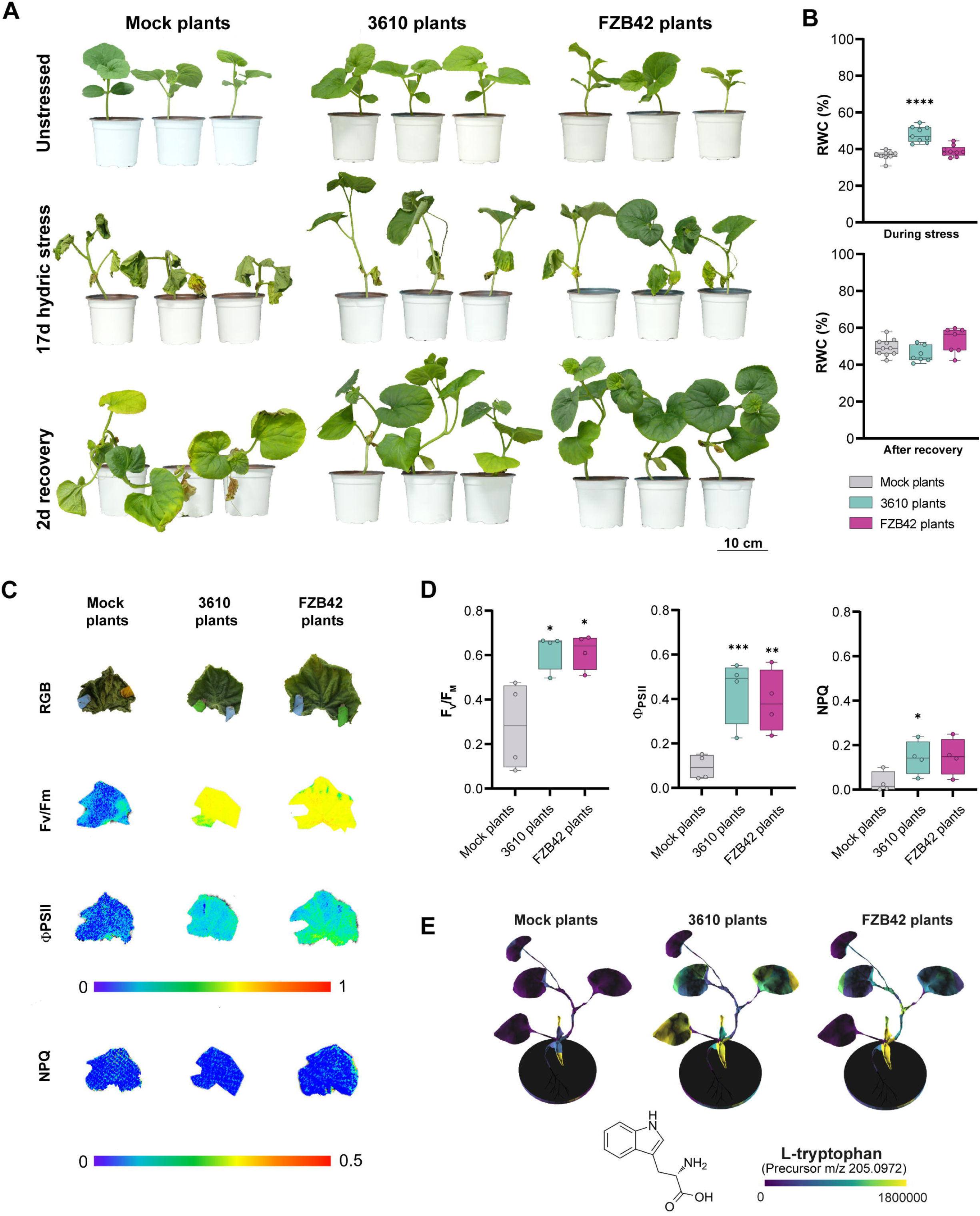
*Bacillus subtilis* seed priming enhances drought resilience in melon plants. **a** Representative image of mock-, *B. subtilis*-, and *B. velezensis*- primed plants under three conditions: well-watered (top), after 17 days of drought (middle), and 2 days post-rewatering (bottom). **b** Relative water content (RWC) of leaves from mock- and *Bacillus*-treated plants under drought and after rewatering. Box and whisker plot. One-way ANOVA with Dunnett’s test (****P < 0.0001 vs. mock). **c** RGB images, F_V_/F_M_, Φ_PSII_ (steady-state), and NPQ (steady-state) measured 2 days after rewatering in leaves from each treatment group. **d** Quantitative values of F_V_/F_M_ (left), Φ_PSII_ (middle), and NPQ (right). One-way ANOVA with Dunnett’s test (*P = 0.0134; **P = 0.0017; ***P = 0.0003). **e** Spatial distribution of L-tryptophan abundance (spectral match) in 3D models of mock- and *Bacillus*-treated plants.

### Species-specific priming against biotic stressors in adult plants

We have previously noted that fengycin-treated seeds confer immunity to the adult plant against the aboveground pathogen *Botrytis cinerea*^13^. To determine whether this response is conserved among *Bacillus* species, we conducted infection assays by inoculating the second leaf of all plant groups with a spore suspension of this phytopathogen. The size of necrotic lesions induced by the fungus in 360 plants was significantly smaller than that of mock or FZB42 plants 3 days post inoculation (d.p.i.). However, by the end of the experiment, at 8 dpi, the size of the necrotic lesions was significantly smaller in both 3610 and FZB42 plants (Fig. 5A). To investigate the physiological underpinnings of this response, we analyzed leaf transpiration patterns following fungal infection by thermography imaging, as they are shown to be reduced in *B. cinerea*-infected melon plants^30^ (Fig. 5B). *B. cinerea* altered whole-leaf transpiration patterns across all treatments compared to non-infected leaves by 3 dpi, based on the reduction on average leaf temperature (Fig. 5B; Fig. S5C,D). Temperature decreases were most pronounced in mock plants (-2.2 °C), followed by FZB42 plants (-1.7°C) and 3610 plants (-1.4°C). By 8 dpi, leaf temperatures returned to pre-infection levels in all groups except FZB42 plants, whose temperature exceeded those values. To assess localized effects of *B. cinerea* infection and its spatial spread, transects (∼12 mm; 35 pixels) centered on the inoculation sites were analyzed from thermal images. Temperature values for each pixel have been plotted on the corresponding profiles (Fig. 5B). At 3 dpi, mock plants displayed the lowest temperatures across transects, particularly at the inoculation point. By 8 dpi, overall transect temperatures increased, with the most pronounced rise occurring at the inoculation point. In contrast, 3610 plants exhibited higher, more uniform temperatures across transects at 3 dpi, with minimal fluctuations at 8 dpi. It is worth noting that 3610 plants at 3 dpi reached temperature profiles similar to those of mock plants at 8 dpi, suggesting an earlier detection of the pathogen. FZB42 plants, however, showed an early localized temperature increase at 3 dpi at the inoculation site, while surrounding areas remained comparable to mock plants. By 8 dpi, FZB42 plants exhibited the highest and most uniform temperatures across the analyzed transect, with no apparent temperature gradients.

**Figure 5.**
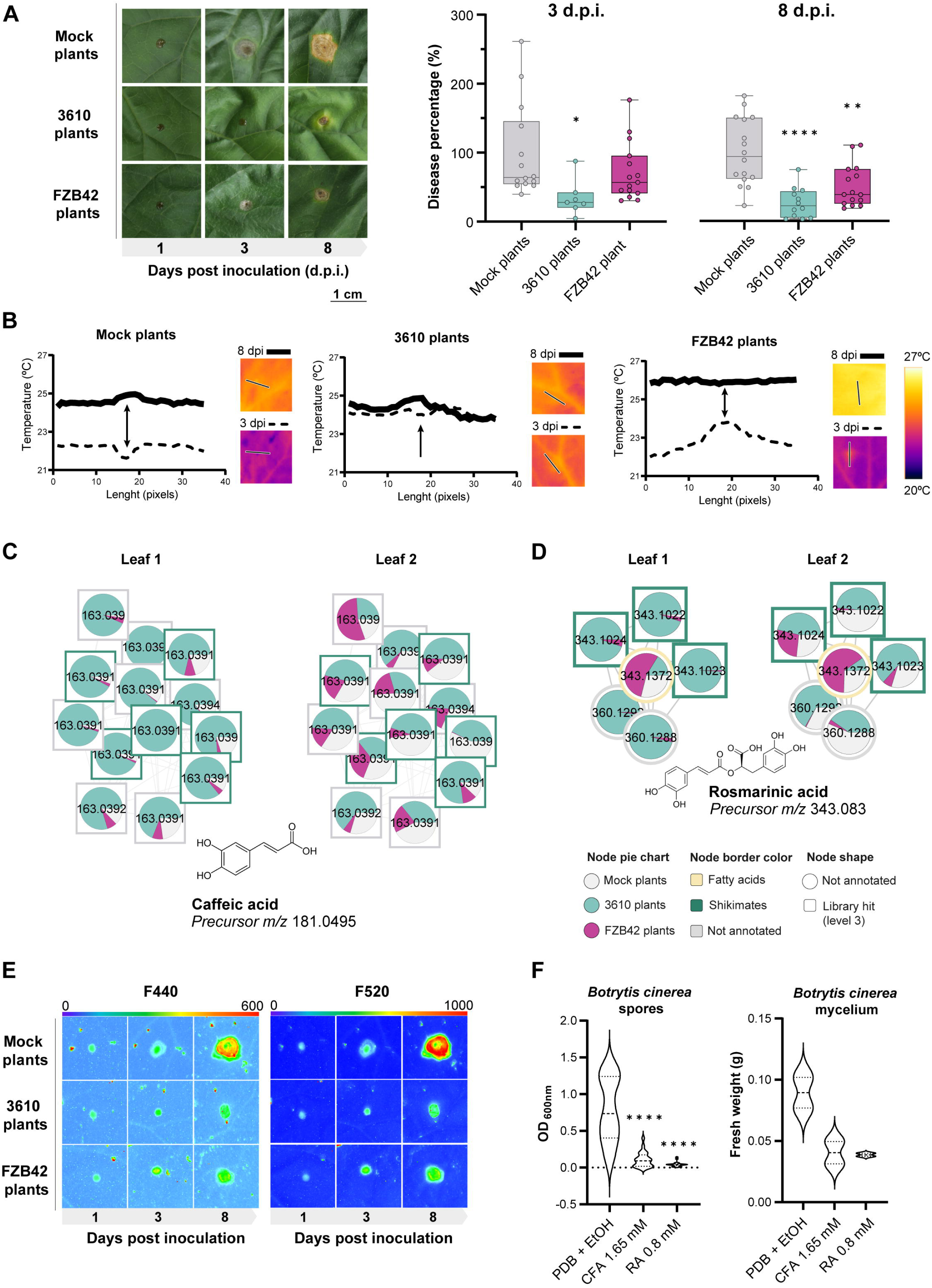
Seed priming with *Bacillus* strains enhances phenolic accumulation and resistance to *Botrytis cinerea* in melon. **a** Left: Necrotic lesions in leaves from mock-, *B. subtilis*-, and *B. velezensis*-treated plants, 1–8 days after *B. cinerea* inoculation. Right: Lesion area quantification normalized to mock-treated controls at 3 and 8 dpi. Box-and-whisker plots. One-way ANOVA with Dunnett’s test (*P = 0.0198; **P = 0.0012; ****P < 0.0001). **b** Leaf surface temperature profiles (35-pixel transect) across lesions and surrounding tissue. Central inoculation point indicated with an arrow. **c, d** Molecular family networks of caffeic acid (CFA) (c) and rosmarinic acid (RA) (d) and related phenolics differentially accumulated in leaves from mock- and *Bacillus*-treated plants. Pie charts show mean relative abundance. Node shape: identification level(Sumner et al., 2007); border color: chemical class (NPC#pathway). **e** Fluorescence imaging (F440 and F520) of cell wall–bound phenolics at the inoculation site and adjacent areas. **f** Left: Growth inhibition of *B. cinerea* spores treated with CFA or RA *in vitro* (48-well assay, 3 days). Right: Mycelial growth inhibition with CFA and RA (24 h). Control = PDB + 0.8% ethanol. One-way ANOVA with Dunnett’s test (****P < 0.0001).

To further characterize *B. cinerea* infection dynamics in 3610 and FZB42 plants, where disease percentage was significantly reduced, specific metabolites potentially involved in defense responses were analyzed. Non-targeted metabolomics identified significant accumulation of caffeic acid (CFA) and rosmarinic acid (RA; an ester of caffeic acid and 3,4-dihidroxyphenyllactic acid) in older leaves of 3610 plants (Fig. 5C,D; Table S2). These metabolites may localize within the apoplast or become esterified to the cell wall. To clarify their anatomical distribution, melon plants were illuminated with UV light, and the fluorescence emitted by phenolic compounds covalently bonded^31^ to the cell wall was recorded in the blue (F440) and green (F520) regions of the spectrum. Blue-Green Fluorescence (BGF) emission was measured in the second leaf of adult-stage healthy melon plants (Fig. S6A,B). No significant differences were observed in F440 and F520 emission either in the spatial pattern or averaged over the whole leaf across the three treatments in the absence of pathogen inoculation. On the other hand, BGF emission of *B. cinerea*-inoculated leaves was also recorded to test whether the compounds remained soluble or esterified by enhancing the known increase in F440 and F520 emissions of the affected areas^30^ (Fig. 5E). At 3 dpi, fungal lesions on mock and FZB42 plants could be visualized as a spot with higher F440 and F520 emission than the surrounding areas. Interestingly, in 3610 plants, the *B. cinerea*-affected area did not increase in size at 3 dpi. By the final measurement, the affected area expanded in all treatments, as indicated by higher F440 and F520 emissions. Mock plants showed the highest F440 and F520 emissions and the largest affected area, followed by FZB42 plants and finally 3610 plants, which showed the lowest BGF emission and the narrowest affected area. When *B. cinerea* spores were inoculated with commercial caffeic acid and rosmarinic acid, their growth was inhibited compared to non-treated spores. Similarly, treatment of already germinated spores with these compounds arrested further mycelial growth (Fig. 5F). Interestingly, these compounds did not impair growth or biofilm formation of either *B. subtilis* or *B. velezensis* when tested *in vitro* (Fig. S6C,D). These findings indicate that soluble phenolic compounds accumulated in leaves due to *Bacillus* treatment of the seeds are cytotoxic on *B. cinerea* likely preventing fungal spread to surrounding areas and, thus, reducing necrotic lesion size.

Previous studies have associated the accumulation of plant specialized metabolites, including terpenoids, green-leaf volatiles and aromatic compounds to defense against herbivores^32^. While terpenoids play a crucial role in attracting natural enemies both above- and below-ground^33^, alkaloids are well known for their potent toxicity against various organisms, including arthropods, by interacting with DNA, membranes and enzymes^34^. In addition, chlorogenic acid has been described as an important metabolite in defense against herbivores^35^. Given the accumulation of these metabolites in plants grown from *Bacillus*- treated seeds (Fig. 2C; Fig. S4A,B), we conducted experiments to assess whether this observation translates into enhanced herbivore resistance. We examined to what extent the metabolic status of leaves and their putative volatile compound profile influence herbivore attraction or deterrence. To test this, we conducted preference assays on melon leaves using two major agricultural pests, *Bemisia tabaci* (whitefly) and two strains of *Tetranychus urticae* (two-spotted spider mite)^36^. After a maximum of 15 minutes of free flight, whiteflies landed more frequently on mock plants than on those grown from *B. subtilis*-treated seeds. This behavior was not observed when mock plants were faced with FZB42 plants or when mites were tested, given that no significant differences in leaf choice were detected in any pairwise comparison (Fig. 6A). However, oviposition assay in non-choice condition showed a similar reproductive performance in all treatments. In contrast, mite reproductive performance was significantly reduced on plants grown from *B. velezensis*-treated seeds in a mite strain previously described as susceptible to induced defenses (*T. urticae* Santpoort-2)^37^ (Fig. 6B). To elucidate the role of the main induced defense pathways i.e. salicylic and jasmonic acid related pathways in this interaction, we performed a gene expression analysis of mite-infested leaves using quantitative reverse-transcriptase PCR (qRT-PCR). Plants grown from seeds treated with *B. velezensis* exhibited upregulation of salicylic acid (SA)-related genes (PR1-1a)^38^ in the absence of herbivore infestation, while JA-related genes (AOS and AOC)^39^ were induced upon infestation, thus, downregulating PR1-1a, probably via the crosstalk between JA and SA pathways^40^ (Fig. 6C). These findings suggest that seed treatment with *B. subtilis* and *B. velezensis* differentially modulates plant- herbivore interactions. *B. subtilis* seed treatment generates plants less attractive to whiteflies, potentially reducing their selection as hosts in natural conditions. In contrast, the above-mentioned initial oxidative burst and JA-mediated defense activation in plants grown from *B. velezensis*-treated seeds, may provide protection against mite infestation during later developmental stages.

**Figure 6.**
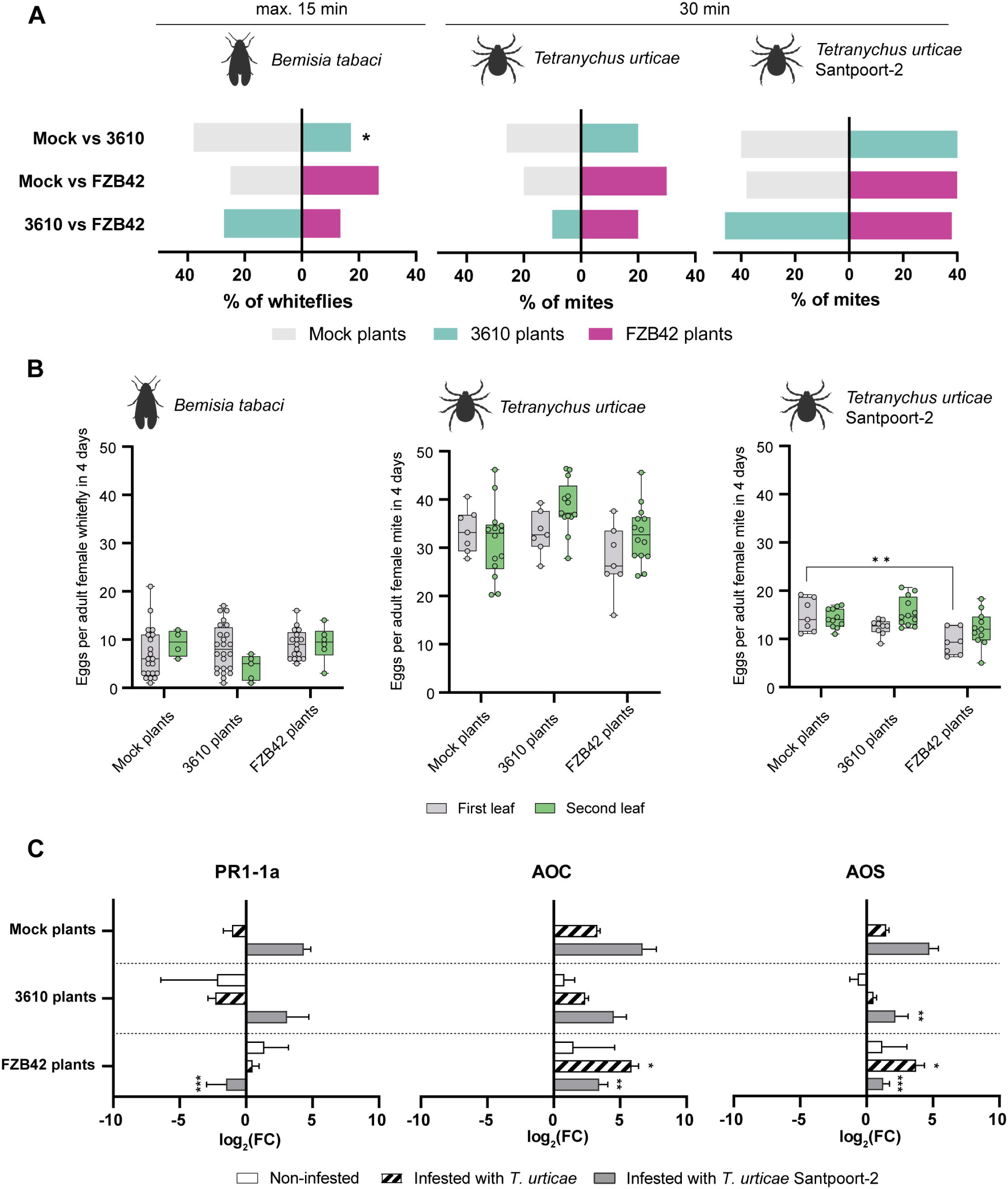
Seed priming with *Bacillus* alters plant–herbivore interactions, reducing whitefly attractiveness and impairing performance of JA-sensitive spider mites. **a** Choice bioassays showing preference of *Bemisia tabaci* whiteflies, *Tetranychus urticae*, and the Santpoort-2 strain of *T. urticae* for melon leaves from mock-, *B. subtilis*-, and *B. velezensis*-treated plants. In each pairwise comparison, the percentage of individuals selecting each treatment is shown. Time points are indicated above graphs. Statistical significance: chi-squared test (*P = 0.0339; n = 49–58 per group). **b** Reproductive performance (number of eggs/female in 4 days) of *B. tabaci*, *T. urticae*, and *T. urticae* Santpoort-2 on first and second leaves. Box-and-whisker plots show distribution across replicates (dots = individual values). One-way ANOVA with Dunnett’s test (**P < 0.001 vs. mock). **c** Relative transcript abundance (log₂FC) of defense-related genes in second leaves of mock- and *Bacillus*-treated plants under non-infested or mite-infested conditions. Genes include PR1-1a (SA pathway), AOS and AOC (JA pathway). Values normalized to *actin*-7 and expressed relative to non-infested mock plants. Mean ± SE. Two-way ANOVA with Dunnet’s test (*P < 0.1, **P < 0.001, ***P < 0.0001).

## Discussion

By comparing two phylogenetically related beneficial bacteria applied at the seed stage, we demonstrate that early microbial priming leads to distinct physiological and metabolic programs in *Cucumis melo*. Despite similar colonization efficiency and persistence, *B. subtilis* and *B. velezensis* induced divergent developmental dynamics, leaf metabolite profiles, and stress-related responses. These results underscore the plant’s capacity to functionally differentiate between beneficial microbial partners and tailor its physiological trajectory accordingly.

Notably, *B. velezensis* induced a transient delay in radicle emergence, particularly at higher doses, suggesting a threshold-dependent effect on early growth. Previous work has shown that structural variants of lipopeptides produced by *Bacillus* strains can modulate biological activity and host perception, potentially accounting for the delay observed^41^. Despite early modulation of growth, plants treated with *B. velezensis* reached developmental stages comparable to controls, indicating that early reprogramming does not compromise long-term performance, in line with current paradigm highlighting the existence of a recovery phase in which plants either reset or consolidate stress memory^42^. Transcriptomic data revealed upregulation of retrotransposon activity and repression of proteasome-related genes in seeds treated with *B. velezensis*, suggesting a regulatory state in which genomic flexibility is increased while proteolytic activity is restrained. Such regulation may reflect early adjustments in developmental plasticity. While our data do not directly address epigenetic mechanisms, parallels with stress memory upon pathogenic infection^43,44^ and mutualistic fungal interactions^2,45^ raise compelling questions about the role of chromatin-level reprogramming in bacterial priming.

At the whole-plant level, *Bacillus*-treated plants displayed species-specific metabolite accumulation patterns in leaves. While both strains maintained photosynthetic efficiency, they differed in stress resilience. *B. subtilis* enhanced drought tolerance, as shown by increased relative water content, starch accumulation in chloroplasts, and elevated levels of L-tryptophan, previously linked to hydric stress responses^29^. Conversely, *B. velezensis* induced an intermediate phenotype under water-limited conditions, marked by some accumulation of L-tryptophan and increased levels of Luteolin-7-glucoside^46^. Specialized metabolites with antifungal properties, including caffeic acid, rosmarinic acid, and a chlorogenic acid analog, were enriched in leaves of *Bacillus*-primed plants. This metabolic reconfiguration likely contributes to enhanced resistance against *Botrytis cinerea*, as observed in both treatments. In addition, priming effects extended to herbivore interactions: *B. subtilis*-treated plants showed reduced attractiveness to whiteflies, while *B. velezensis* treatment led to downregulation of a key jasmonic acid biosynthesis gene in seeds. This may enhance the efficiency of JA-mediated defenses upon attack by spider mites, specifically those populations sensitive to this pathway (Fig. 7). The differential performance of the two *T. urticae* strains tested likely reflects intraspecific variation in herbivore strategies for coping with plant defenses^47^. The Algarrobo- 2 strain, known for its broad host range^48^ and JA-resistance traits^37^, maintained high performance on *Bacillus*-treated plants despite metabolomic shifts, possibly due to enhanced detoxification mechanisms^36^ and the ability to degrade phenolic compounds^49^. In contrast, the Santpoort-2 strain, previously shown to be JA- sensitive^37,50^, was more susceptible to defenses triggered by *B. velezensis* seed treatment, supporting the role of JA pathway priming in herbivore resistance. Whiteflies exhibited a reduced preference for *B. subtilis*-treated plants, although their performance was not affected. While this suggests potential interference with pest establishment, further studies under field-like conditions are needed to assess long-term population dynamics. Altogether, these findings indicate that early microbial priming can alter plant-herbivore interactions in a strain-specific manner, offering protective effects against key pests. Future work should explore the bidirectional influence between microbial colonization and herbivore-induced defenses, which may shape both microbiome composition and plant resistance traits over time^51,52^. Certainly, although our work focused on host responses, the inoculation of seeds with *Bacillus* strains likely reshapes the seed and root microbiome. Early microbial colonizers are known to influence the assembly of microbial communities during plant development^53,54^. Further studies will be required to assess how this restructuring affects long-term plant–microbiome dynamics and contributes to the phenotypes observed in this study.

**Figure 7.**
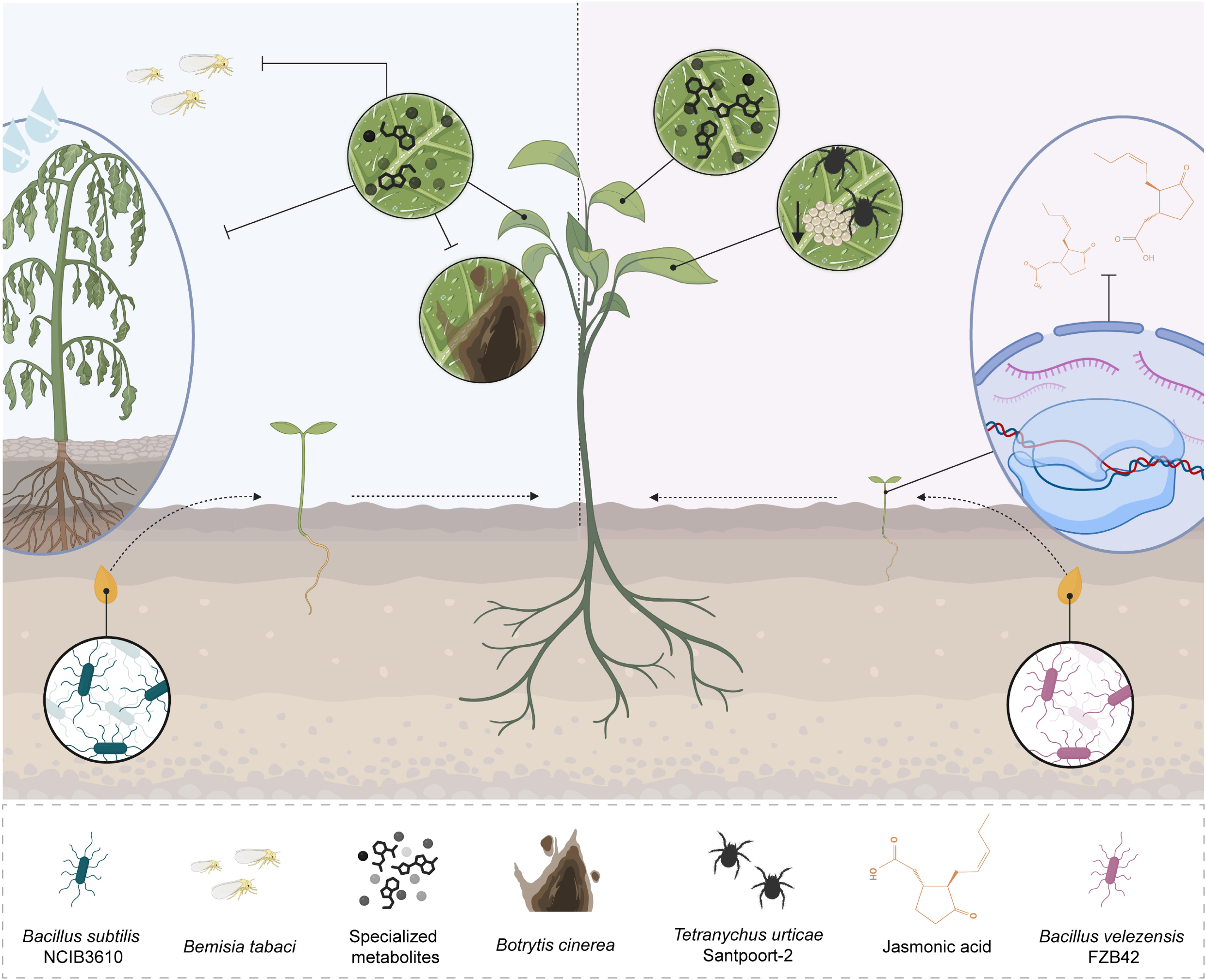
Schematic representation of distinct plant programs triggered by seed priming with *Bacillus subtilis* (left) and *Bacillus velezensis* (right). Overview of key phenotypic, metabolic, and defensive outcomes in melon plants following early priming with each strain. Created with BioRender.com.

In conclusion, our findings reveal that even among closely related mutualistic microbes, early interactions can lead to divergent physiological programs. These results challenge the assumption of functional redundancy among beneficial strains and highlight the central role of the host in modulating mutualistic outcomes, providing new insight into the diversification and plasticity of beneficial plant–microbe interactions.

## Materials and Methods

### Strains, plant materials and growth conditions

*Bacillus subtilis* NCIB3610 (laboratory collection) and *Bacillus velezensis* FZB42 (laboratory collection) were grown at 28°C from frozen stocks on lysogeny broth (LB: 1% tryptone, 0.5% yeast extract and 0.5% NaCl) plates. The necrotrophic fungus *Botrytis cinerea* (isolate B05.10) was grown at 25°C from a frozen stock in potato dextrose agar (PDA, Oxoid, Thermo Fisher Scientific, Waltham, Massachusetts, USA) plates and maintained until inoculum preparation. Melon seeds (*Cucumis melo* var. Rochet Panal; Semillas Fitó, Barcelona, Spain) were placed in Petri dishes containing wet sterilized filter paper, covered with foil, and kept in growth chambers at 25 °C for 5 days when studying short-term effects. For long-term growth promotion evaluation and further analysis, seedlings from treated seeds were transferred to pots and maintained in a growth chamber (25°C, 16L/8D photoperiod, 60% relative humidity, 300 µmol/s/m^2^). *Arabidopsis thaliana* ecotype Columbia-0 (Col-0) seeds were grown in Murashige and Skoog medium plates after 2-day stratification in darkness at 4°C upon radicle analysis.

### Herbivores

*Tetranychus urticae* Algarrobo-2 (La Mayora, Spain) is a red form originally collected from field visits in Málaga (Spain), and shows that can infest bean, melon and tomato. *T. urticae* Santpoort-2 colonies were provided by Dr Merijn Kant and were originally collected from *Euonymus europaeu*s (spindle tree) located in the Natural Reserve of Santpoort. This colony has been described as susceptible to tomato induced defenses^37,55^. Mite colonies were reared on detached leaves of *Phaseolus vulgaris* cv. Canadian Wonder, placed on wet cotton wool and maintained in a climate room (25°C, 16L/8D photoperiod). 16 days old females were used for the bioassays collected from eggwaves. Briefly, adult females were placed on a clean bean leaf and removed after 48 h. Colonies were maintained in the same conditions as described above for 16 days. *Bemisia tabaci* Mediterranean (MED) individuals were obtained from a colony originating from individuals collected during field visits in Málaga (Spain) and reared on melon plants (*Cucumis melo* L. ‘ANC42’, IHSM seedbank collection, La Mayora) in wooden cages covered with insect-proof nets, in an insect-proof glasshouse with temperature control (22 to 27°C day and 17 to 20°C night) and light supplementation when needed as described in ref^56^. We used 3 ± 1 days old adult females for the assays. To obtain same age females, we used a colony from the collection and remove all adult whiteflies with an aspirator. We used the newly emerged females 24h after removing the adults.

### Quantitative analysis of radicle and plant growth

Seeds were *Bacillus*-inoculated as previously described^13^. To obtain cell suspensions, cultures grown in liquid LB at 28°C overnight were washed twice with distilled water and adjusted to the specified optical density (OD_600_) corresponding to 10^7^ CFU/ml (lower inoculum) or 10^8^ CFU/ml (higher inoculum). Seeds were surface-sterilized with 0.15% sodium hypochlorite and treated with bacterial cell suspensions by immersion for 1 hour and 30 minutes at room temperature with mild shaking. Seeds treated with water were used as controls and considered mock treatment. To assess radicle growth-promoting effects, radicle areas were measured 5 days post-treatment using ImageJ software^57^ (https://imagej.net/ij/). Average growth in the mock group was set to 100% as a baseline to normalize the growth-promoting effects of different bacterial strains. Treatments using specialized metabolites purified from the bacterial cultures were conducted similarly. Fengycin, surfactin and bacillomycin D were purified as described below. At different time points, fresh weight was measured and average growth in the mock group was set to 100% as a baseline to normalize the growth of the groups under study.

### Quantitative analysis of bacterial ecology

For bacterial quantification after treatments, radicles were separated from the seeds and placed in 1.5 ml microcentrifuge tubes containing 1 ml of phosphate-buffered saline (PBS) medium and 1 g of 3-mm glass beads. Attached bacteria were extracted by vortexing the tubes for 1 minute. Seeds were ground in 1 ml of PBS using a mortar and pestle. Total CFU/ml were assessed by plating serial dilutions of the suspensions on solid LB medium. To quantify spores, the same suspensions were subjected to a 5-minute thermal shock at 80 °C before plating. For bacterial localization on roots, *B. subtilis* and *B. velezensis* strains resistant to spectinomycin and chloramphenicol, respectively, were used for seed inoculations. Roots were collected in 10 ml of distilled water and homogenized for 3 min using a stomacher-type homogenizer. Total CFU/ml and spore CFU/ml were assessed by plating serial dilutions of the suspensions on solid LB medium supplemented with the appropriate antibiotic for strain selection.

### RNA sequencing

Total RNA was extracted of each condition from two different replicates using RNAprep Pure Tissue Kit (TIANGEN). cDNA library preparation and RNA sequencing were performed by Novogene using an Illumina NovaSeq 6000 or NovaseqX plus sequencers. For LncRNA, firstly, ribosomal RNA was removed by rRNA removal kit (TruSeq Stranded Total RNA Library Prep), and rRNA free residue was cleaned up by ethanol precipitation. Subsequently, sequencing libraries were generated using the rRNA-depleted RNA using NEBNext® Ultra™ Directional RNA Library Prep Kit. After fragmentation, the first strand cDNA was synthesized using random hexamer primers. Then the second strand cDNA was synthesized and dUTPs were replaced with dTTPs in the reaction buffer. The directional library was ready after end repair, A-tailing, adapter ligation, size selection, USER enzyme digestion, amplification, and purification. For smallRNA, 3’ and 5’ adaptors were ligated to 3’ and 5’ end of small RNA, respectively. Then the first strand cDNA was synthesized after hybridization with reverse transcription primer. The double-stranded cDNA library was generated through PCR enrichment. After purification and size selection, libraries with insertions between 18∼40 bp were ready for sequencing on Illumina sequencing with SE50. Both libraries were checked with Qubit 3.0 and real-time PCR for quantification and an Agilent 2100 bioanalyzer for size distribution detection. Quantified libraries were pooled and sequenced on Illumina platform NovaseqX plus (LncRNA, mRNA and other ncRNA) and Novaseq6000 (smallRNA), according to effective library concentration and data amount. The clustering of the index-coded samples was performed according to the manufacturer’s instructions. After cluster generation, the library preparations were sequenced and paired-end reads were generated (strategy PE150) with 12 Gb (40M reads) output per sample in the case of LncRNA and in single-end read (strategy SE50) with 10M reads output/ sample in the case of smallRNA. Differential expression gene analysis from Novogene with reference genome (*Cucumis melo cv.* DHL92 v4)^58^ was further evaluated.

### Lipopeptides extraction and purification

Lipopeptides of interest were obtained by incubating a liquid culture of *Bacillus velezensis* FZB42 for 48 hours and centrifuged (25 min, 8500 *g*, 4°C). The supernatant, containing lipopeptides, was collected. Protein and peptide precipitation was induced by adjusting the supernatant to pH 2 with 6N HCl, and the solution was incubated overnight at 4°C. The precipitate was collected by centrifugation (25 min, 8500 *g*, 4°C) following pellet extraction with methanol under agitation overnight. Insoluble material was removed by centrifugation and the supernatant was concentrated by rotary evaporation. The crude lipopeptide extract was purified by solid-phase extraction (SPE) using Strata C18-U cartridges (200 mg, 3 mL column volume (CV), Phenomenex, Torrance, CA, USA). Cartridges were activated with one CV of methanol, conditioned with one CV of ultrapure water (Milli-Q system, Millipore, Billerica, MA, USA) and loaded with the sample. Sequential elutions were performed with Milli-Q water (1 CV) and increasing methanol concentrations in water (25–100%). Fractions from each wash were analyzed by thin layer chromatography (TLC) and high-performance liquid chromatography (HPLC) to identify lipopeptide-containing fractions. For TLC, a mobile phase of dichloromethane (65%), methanol (25%) and water (4%) was used. Each fraction was spotted onto silica gel plates, developed in the mobile phase, and dried. Plates were then sprayed with water to visualize lipopetide presence. Lipopeptides analysis and purification were performed using an Agilent™ 1260 Infinity II HPLC (Agilent, Santa Clara, CA, USA) with a gradient method of a mobile phase of acetonitrile and 0.1% trifluoroacetic acid (TFA) in Milli-Q water following a UV-visible detection method at 210 nm. The method included an analytical phase with a 250 x 4.6 mm, 5 µm Zorbax Eclipse Plus C18 column (Agilent) with 20 µL injections of each SPE fraction, followed by a preparative phase on a 250 x 9.8 mm, 5 µm Zorbax Eclipse Plus C18 column (Agilent) with successive 350 µL injections of the 75% and 100% methanol SPE extracts for final lipopeptide purification. Lipopeptides were isolated based on retention times from the analytical phase (40/60 for bacillomycin D, 50/50 for fengycin and 80/20 for surfactin) and concentrated under nitrogen flow. To confirm chromatographic peaks as bacillomycin D, fengycin or surfactin, aliquots from these fractions were analyzed by Matrix-Assisted Laser Desorption/Ionization Time of Flight (MALDI-TOF) mass spectrometry (UltrafleXtreme Bruker, Billerica, MA, USA). The results were compared to known mass/charge (m/z) values for each lipopeptide.

### Metabolite extraction from plant tissues

Adult melon plants derived from both treated and untreated seeds were sectioned to analyze the entire plant metabolome. The roots, stems, and leaves were each divided into multiple parts (11 sections from roots, 15 from stems, and 3–4 sections from each leaf). Metabolites were extracted from the samples by adding 1 ml of 80% HPLC-grade methanol, followed by vigorous vortexing. Only one replicate was performed because of the high number of sections analyzed per plant. Given the large number of samples, weighing and normalization by biomass were not feasible; however, the size of each plant section was kept constant. To allow for relative quantitative comparisons between samples, peak areas were normalized to total ion count, a robust and commonly used strategy in non-targeted metabolomics^59^. Additionally, sulfamethazine (1 µM) was included in all samples as an internal standard and used for normalization during analysis. Following a 2-hour incubation, extracts were centrifuged at 14,000 g for 30 minutes. The resulting supernatants were stored at -80°C until analysis by liquid chromatography-tandem mass spectrometry (LC-MS/MS). Extraction controls were prepared by processing three blank samples containing only pure methanol. Metabolite abundance differences were estimated based on relative comparisons among equally treated samples, as authentic standards were not available in the non-targeted metabolomics approach.

### Liquid chromatography-tandem mass spectrometry

Non-targeted metabolomics was carried out using LC-MS/MS on a Q-Exactive Quadrupole-Orbitrap mass spectrometer coupled to a Vanquish ultra-high-performance liquid chromatography (UHPLC) system (Thermo Fisher Scientific), following the protocol described in ref^60^. A 5 μL aliquot of each sample was injected into a UHPLC system equipped with a Kinetex C18 core-shell column (50 × 2 mm, 1.8 μm particle size, 100 Å pore size; Phenomenex, Torrance, USA).

Chromatographic separation was conducted at a flow rate of 0.5 mL/min using solvent A (H₂O with 0.1% formic acid) and solvent B (acetonitrile with 0.1% formic acid) as the mobile phases. The linear gradient was applied as follows: 0–0.5 min, 5% solvent B; 0.5–4 min, 5–50% solvent B; 4–5 min, 50–99% solvent B; 5– 7 min, washout at 99% solvent B; and 7–9 min, re-equilibration at 5% solvent B. In positive ion mode, electrospray ionization (ESI) was used with the following settings: sheath gas flow of 35 AU, auxiliary gas flow of 10 AU, sweep gas flow of 2 AU, and auxiliary gas temperature of 400 °C. The spray voltage was set at 3.5 kV, the inlet capillary temperature at 250 °C, and the S-lens voltage at 50 V. MS/MS spectra were obtained using data-dependent acquisition (DDA) mode. Full MS1 scans (m/z 150–1,500) were recorded at a resolution of 17,500, followed by up to five MS/MS scans per duty cycle. The C-trap fill time was limited to a maximum of 100 ms or until the automatic gain control (AGC) target of 5 × 10⁵ ions was reached. Precursor ions were isolated with a quadrupole window of 1 m/z. Fragmentation was carried out using normalized collision energies (NCE) of 20%, 30%, and 40%, with z = 1 as the default charge state. Apex triggering was applied to acquire MS/MS scans 2–15 seconds after the precursor’s initial detection in the MS1 survey scan. A dynamic exclusion window of 5 seconds was used. Precursor ions with unassigned charge states and isotope peaks were excluded from MS/MS analysis.

### Feature-based molecular networking and spectral library search

Following LC-MS/MS data acquisition, MS1 and MS/MS feature extraction was conducted using MZmine software version 3.9.0^61^. Peak detection thresholds were set at 1E5 for MS1 and 1E3 for MS/MS. For building MS1 chromatograms, a mass accuracy of 10 ppm and a minimum peak intensity of 5E5 were applied. Extracted Ion Chromatograms (XICs) were deconvoluted using the baseline cutoff algorithm with a minimum intensity threshold of 1E5. After deconvolution, XICs were matched to MS/MS spectra using a 0.02 m/z and 0.2-minute retention time window. Isotopic peaks were grouped, and features across different samples were aligned with a mass tolerance of 10 ppm and a retention time tolerance of 0.1 min. MS1 features lacking associated MS/MS spectra were removed from the dataset, along with those that did not include isotope peaks or were present in fewer than three samples. Gaps in the resulting feature matrix were filled using a relaxed retention time tolerance of 0.2 min and a mass tolerance of 10 ppm. The final feature table was exported as a .csv file, and the corresponding MS/MS spectra were saved in .mgf format. Contaminant features detected in blank samples were filtered out; only those with a blank-to-average relative abundance ratio below 30% were retained for further analysis. For feature-based molecular networking and spectral library matching, the .mgf file was uploaded to GNPS (https://gnps.ucsd.edu/ProteoSAFe/static/gnps-splash.jsp)^62^. Molecular networking parameters included a minimum cosine score of 0.7, precursor ion mass tolerance of 0.01 Da, fragment ion mass tolerance of 0.01 Da, minimum of 4 matched fragment peaks, and a minimum cluster size of 1 (with MS Cluster deactivated). For analog searches, the same cosine threshold (0.7) was used, with a maximum allowed analog search mass difference of 100 m/z. Networks were visualized using Cytoscape software version 3.10.2^63^. To enrich chemical structural annotation in the molecular networks, in silico predictions were integrated using SIRIUS 4 software (https://bio.informatik.uni-jena.de/software/sirius/)^64^, comparing input data against internal databases such as Bio Database, GNPS, Natural Products, PubChem, and PubMed. Chemical classes were annotated using NPClassifier^65^ and ClassyFire ^66^ ontologies. Mirror plots comparing the *mzspec* values of selected features with database records were generated using the USI resolver tool (https://metabolomics-usi.ucsd.edu/) (Fig. S8). Annotations followed the guidelines outlined in ref^67^ (Tables S2 and S3). Statistical analyses were conducted in Metaboanalyst v.5.0^68^ (https://www.metaboanalyst.ca/) using the quantification table after filtering by interquartile range (IQR) and normalizing based on the internal standard sulfamethazine.

### 3D mass spectral molecular cartography and visualization

Collection of plant material for 3D mass spectral molecular cartography and 3D modelling was done as described in ref^13^. Point coordinates and LC-MS data were combined into a .csv file and mapped on 3D models using Ili software ^69^ (https://ili.ucsd.edu/).

### Hydric stress assay

Hydric stress assays were conducted in 21-day-old plants grown from untreated seeds and seeds treated with *B. subtilis* and *B. velezensis*. Under normal conditions, the plants were irrigated with 1.5 L of water per tray containing 16 plants, twice a week. For the hydric stress assay, irrigation was withheld for 17 days. Recovery was assessed by taking measurements two days after rewatering. Relative water content (RWC) of leaves was recorded before, during and after hydric stress period. Six samples were collected per treatment, with 4- 5 leaf discs (1.5 cm of diameter) taken per sample. RWC was calculated using the formula *RWC=(FW-DW)/(TW-DW)*, where FW is the fresh weight of the samples, TW is the turgid weight measured after hydration for 3-4 h at RT, and DW is the dry weight obtained after oven-drying the samples at 80°C for 24 h. RWC was expressed as a percentage.

### Analysis of photosynthetic capacity

The photosynthetic performance of plants grown from mock-treated seeds, as well as from seeds treated with *B. subtilis* and *B. velezensis*, all under non-stressed and water-stressed conditions, was evaluated by monitoring PSII activity by variable chlorophyll fluorescence imaging (Chl-FI) using an Open FluorCam 700 MF (Photon System Instruments, Brno, Czechia). Thirty minutes dark-adapted leaves were placed below the camera and measured according to the Protocol 1 described in ref^70^. Firstly, F_0_ was recorded. Then, a short saturating light pulse of 2500 µmol photons/m^2^·s was applied and F_M_ was measured; subsequently, the leaf was illuminated with 400 µmol photons/m^2^·s of photosynthetic active radiation light and eleven successive saturating light pulses were applied to measure F_t_ and F’_M_ at 2, 5, 10, 20, 30, 60, 90, 120, 180, 240 and 300 s. Using FluorCam software version 5.0 (Photon System Instruments), black-and-white images of transient chlorophyll fluorescence were collected and used to calculate several fluorescence parameters, according to ref^71^: a) maximum quantum yield of PSII [F_v_/F_M_= (F_M_-F_0_)/F_M_]; b) effective quantum yield of PSII (Φ_PSII_= (F’_M_-F_0_)/F’_M_); and c) non-photochemical quenching [NPQ = (F_M_-F’_M_)/F’_M_]. A false-color scale was then applied to these calculated images in the FluorCam software to visualize parameter variations. The images (512x512 pixels resolution) presented in figures correspond to the most representative data.

### Transmission electron microscopy (TEM)

Leaves of adult plants were split and cut into sections of 0.5 cm x 0.5 cm approximately. Samples were fixed directly using 4% (v/v) paraformaldehyde and 2.5% (v/v) glutaraldehyde in 0.1 M phosphate buffer (PB) overnight at 4°C. Following three washes in PB, the sections were post-fixed in a 1% osmium tetroxide solution in PB for 90 min at room temperature. Subsequently, samples underwent PB washes and 15 min of stepwise dehydration in an ethanol series (30%, 50%, 70%, 90% and twice in 100%). Between the 50% and 70% ethanol steps, the samples were incubated in-bloc in 2% uranyl acetate solution in 50% ethanol at 4°C overnight. After dehydration, samples were progressively embedded in LR White resin as follows: resin:ethanol (1:1) for 4h, resin:ethanol (3:1) for 4h and pure resin overnight. The sample blocks were embedded in capsule moulds containing pure resin for 72h at 70°C. The samples were left to dry and visualized under a FEI TALOS F200X (Thermo Fisher Scientific). To estimate starch granule size (µm^2^), free-hand selection of granules of at least five different sections within three replicates of each plant group were analyzed using ImageJ software.

### Scanning electron microscopy (SEM)

For SEM analysis, leaves of adult plants were split and cut in sections of 0.5 cm x 0.5 cm approximately. Samples were fixed directly using 2.5% (v/v) glutaraldehyde in 0.1 M PB overnight at 4°C. After three washes in PB, samples were dehydrated in an ethanol series (50%, 60%, 70%, 80%, 90% and 100% twice) and critical point drying was performed. Total number of stomata was counted from at least five different sections within three replicates of each plant group and assessed as stomatal density (S/mm^2^) by dividing by area.

### Quantification of chlorophyll *a* and chlorophyll *b*

Quantification of chlorophylls was performed according to ref^72^. Briefly, 3 leaf disks (1.5 cm of diameter) per plant were collected from ten plants in each group. Chlorophyll was extracted using 80% acetone, followed by centrifugation at 9000 *g* for 5 min, and absorbance was measured at 647 nm and 663 nm on a spectrophotometer. Chlorophyll concentrations were calculated using the equations: *[Chl a] (µg/ml) = 12.25A_663_ – 2.79A_647_* and *[Chl b] (µg/ml) = 21.5A_647_ – 5.10A_663_* ; and expressed as µg/cm^2^.

### Fungal infection assays in plants

*B. cinerea* infection assays were conducted in 3-4-week-old plants grown from untreated seeds and seeds treated with *B. subtilis* and *B. velezensis*. Fungal spores were harvested from light-grown culture in sterile distilled water and filtered through a 40 µm cell strainer to remove remaining hyphae. The conidial suspension for inoculation was adjusted to 10^5^ spores per ml in half-strength filtered (0.45 µm) 100% pure organic grape juice. Two distal 5 µl droplet of the conidial suspension were applied to the second leaf of each plant. Plant trays were filled with water, covered with plastic domes and sealed with parafilm to produce high humidity. Leaves were imaged 1, 3 and 8 days post-inoculation and lesion areas were measured using ImageJ software. Lesion sizes were normalized against the average lesion area observed in mock plants, which was set as 100% to represent disease percentage.

### In vitro antifungal activity assays

Cytotoxicity of commercial compounds against *B. cinerea* spores and mycelium growth was assessed by absorbance measurements of fungal biomass at 600 nm (spores stage) and fresh weight measurements (mycelium stage). Compounds were tested in 96-well plates across a range of concentrations, with a final volume of 100 µl per well. Once growth inhibition was observed, experiments with specific concentrations were performed. To assess spore growth, 100 spores per well were inoculated from a fresh conidial suspension and absorbance at 600 nm was measured after 48h. For mycelium biomass, 1 ml of saturated potato dextrose broth (PDB) containing fungal biomass was inoculated per well. Fresh weight was measured after 24h using 0.45 µm filters to remove residual PDB. PDB supplemented with the maximum concentration of ethanol used for compound resuspension was used as control condition for both assays.

### Bacterial growth and biofilm formation

Citotoxicity of commercial compounds against *B. subtilis* and *B. velezensis* was assessed by quantitative analysis of bacterial growth and biofilm formation. To evaluate bacterial growth, commercial compounds were added to 5 ml of LB medium and 10 µl of liquid culture grown overnight. Total CFU/ml were assayed as described above. Biofilm assays were performed on Msgg medium: 100 mM morpholinepropane sulfonic acid (MOPS) (pH 7), 0.5% glycerol, 0.5% glutamate, 5 mM potassium phosphate (pH 7), 50 µg/ml tryptophan, 50 µg/ml phenylalanine, 50 µg/ml threonine, 2 mM MgCl_2_, 700 µM CaCl_2_, 50 µM FeCl_3_, 50 µM MnCl_2_, 2 µM thiamine, 1 µM ZnCl_2_. To analyze biofilm formation under presence of commercial compounds, 10 µl of washed liquid culture were inoculated in 1 ml of liquid Msgg in 48-well plates and incubated at 28°C. Results were observed at 24h.

### Thermal imaging

Infrared images of leaves of plants grown from mock-treated and *Bacillus*-treated seeds, under both uninfected and *B. cinerea*-infected conditions, were taken in the growth chamber using a Photon A305sc camera (FLIR Systems, Wilsonville, OR, USA) vertically positioned above the leaf, according to ref^73^. Thermal images were recorded at midmorning in attached leaves. The camera yields images with a resolution of 320x240 pixels and thermal sensitivity <0.05°C in the spectral range of 7.5-13 mm, and it is operated with FLIR Research & Development software version 3.4, which was also used to obtain temperature values from measurements. Numerical data from 11 regions of interest per plant were analyzed. The images in figures, shown using a false color scale, correspond to standard experiments.

### Blue-green fluorescence imaging (BGF)

Under UV light, phenolic compounds emit fluorescence in the blue (F440) and green (F520) regions of the spectrum that can be recorded by a customized Open FluorCam FC 800-O (Photon Systems Instruments), following the method described in ref^74^. The leaves were placed below the camera and, using an excitation wavelength of 355 nm, nine frames of 800 ms exposure time were taken for each fluorescence wavelength using appropriate cut-off filters. The nine frames were then averaged to compose F440 and F520 black and white images (740×480 pixels resolution). Images of both F440 and F520 were captured from plants grown from mock-treated and *Bacillus*-treated seeds, under both uninfected and *B. cinerea*-infected conditions. FluorCam software version 7.1.0.3 was used to record the images, obtain numerical data from the measurements and apply a false-color scale to the black-and-white images obtained. The images

### Choice and performance assays of *B. tabaci*

Choice assays were conducted using pairwise comparisons of detached leaves from 21-day-old melon plants representing all treatment combinations (mock, *B. subtilis*-treated seeds; *B. velezensis*-treated seeds), following the protocol described in ref^75^ with minor modifications. Briefly, leaves were placed on moist cotton wool inside transparent plastic cages (50 x 50 x 25 cm) for the following comparisons: mock vs 3610, mock vs FZB42 and 3610 vs FZB42. Four leaves (two per treatment) were positioned equidistant to a central flight release platform, alternating between treatments. Female synchronized whiteflies, starved for at least 1 h and briefly chilled on ice, were individually released onto the platform. Plastic cages were placed inside a growth chamber with light intensity at 250 µmol/s/m^2^, 27°C and 70% relative humidity. Only the first choice was recorded; individuals that did not choose within 15 min were excluded. Each whitefly was tested only once, and the cage setup was replaced after every 10 replicates. Fifty individuals were tested per comparison. The distribution of whitefly choice was analyzed using a binomial test against an expected 50:50 ratio for each pairwise assay. Reproductive performance of *B. tabaci* on leaves from plants grown from untreated seeds and seeds treated with *B. subtilis* and *B. velezensis* was assessed following the protocol described in ref^76^. Briefly, three leaf cages were placed in each plant (2 on the first leaf and 1 on the second one), 14 plants per treatment. In each leaf cage, a single adult female whitefly, freshly emerged from colonies maintained separately in the greenhouse, were confined for egg laying. After 4 days, both adults and leaf cages were removed. The leaf portions beneath each leaf cage were observed under stereomicroscope at 40x magnification to count the number of eggs laid. The number of eggs laid per adult female whitefly was used as a measure of reproductive performance.

### Choice and performance assays of *T. urticae*

Choice assays were performed by pairwise comparisons of leaf disks from 21-day-old melon plants grown from untreated seeds and seeds treated with *B. subtilis* and *B. velezensis*. Leaf disks (15 mm in diameter) were placed on wet cotton wool inside a petri dish organized by different pairwise comparisons (mock plants vs 3610 plants, mock plants vs FZB42 plants and 3610 plants vs FZB42 plants). A 2 x 0.5 cm piece of parafilm was used to join the disks in pairs. The plates were rotated and positioned differently within the growth chamber to minimize the influence of external factors on mite choice. A single adult female mite was placed at the center of the parafilm and its choice was recorded after 30 min and 24 h. At least 50 replicates were analyzed for each pairwise comparison. The distribution of mite choices was analyzed using a binomial test against an expected 50:50 ratio for each pairwise assay, pooling all individuals that made a choice. Reproductive performance of *T. urticae* Algarrobo-2 and *T. urticae* Santpoort-2 on leaves from plants grown from untreated seeds and seeds treated with *B. subtilis* and *B. velezensis* was assessed following the protocol described by ^37^. Briefly, five adult female mites were transferred into circular arenas (2-3 cm in diameter) bordered with lanolin (Sigma-Aldrich) onto leaves of 21-day-old melon plants. Three arenas were made per plant (2 on the first leaf and 1 on the second one) and four plants were tested per treatment. After 4 days, the number of eggs was recorded using a stereomicroscope. Spider mite reproductive performance was calculated following the formula *No. of eggs/(live mites + 0.5(total mites – live mites))* for every section. Induction of plant defenses was assessed by measuring the expression of defense-related marker genes. Sections of 21-day-old melon plants emerged from mock-treated and *Bacillus*- treated seeds were infested with adult female spider mites: 15 mites per section, 3 sections per plant (2 on the second leaf and 1 on the first one), 6 plants per treatment. Sections were circular, 2-3 cm in diameter, bordered by lanolin. To assess the basal state of the defense genes in each treatment, we included 6 plants per treatment that were not infested with mites as control plants. At 4 days post infestation (dpi), 6 infested and 6 non-infested plants from each treatment were sampled. Infested sections and corresponding non-infested sections for every group of plants were excised, flash-frozen in liquid nitrogen and stored at -80°C until we carried out mRNA extraction. The three sections obtained from the same plant were pooled to form one biological replicate. Total RNA was extracted from frozen tissue samples using phenol/chloroform extraction with the Macherey-Nagel NucleoSpin RNA Plant kit (Fisher Scientific). For cDNA synthesis, 2 µg of DNase-treated RNA were used, and 1.5 µl of 5-fold diluted cDNA served as the template for 10 µl quantitative reverse-transcription PCR (qRT-PCR) reactions. These were carried out using the SsoAdvanced Universal SYBR Green Supermix (Bio-Rad) on a CFX Opus 384 Real-Time PCR System (Bio-Rad). To assess melon defense responses, the expression of the genes *PR1-1a*, *AOC*, and *AOS* was analyzed (Table S4). Primer efficiency was tested following the method described in ref^77^. Relative transcript levels were calculated using the ΔΔCt method^78^. Gene expression was normalized against the *Actin-7* gene from *Cucumis melo* and expressed as fold-changes in each experimental group (both non-infested and mite-infested plants) relative to the mock-treated, non-infested control group. Each qRT-PCR assay was conducted in triplicate using RNA from six independent extractions.

### Statistical analysis

Statistical differences was assessed by appropriate tests (see figure legends). All analyses were performed using GraphPad Prism version 9 (Boston, Massachusetts USA, www.graphpad.com). P values < 0.05 were considered significant. Asterisks indicate the level of statistical significance: *P<0.05, **P<0.01, ***P<0.001, ****P<0.0001. Statistical analysis of metabolomic datasets was performed by Metaboanalyst v.5.0^68^ (https://www.metaboanalyst.ca/) after filtering by the interquartile range (IQR) and normalizing by feature identified as sulfamethazine. Experiments were repeated at least three times independently with similar results.

## Acknowledgments

We thank Saray Morales for providing technical support, Rocío Camero-Flores for providing support on whitefly management, Cristina Lucena-Serrano, Gregorio Martín-Caballero and Adolfo Martínez-Orellana for technical support on TEM and SEM analysis (SCAI-UMA). This work was supported by grants from ERC Starting Grant (BacBio 637971), Ministerio de Economía y Competitividad I+D+i Plan Nacional (PID2022-141664NB-I00) and Junta de Andalucía, Proyectos de Investigación, Desarrollo e Innovación (I+D+i) (P20_00479). L.C.-R. is funded by a FPI contract (PRE2022-000585) from Ministerio de Ciencia, Innovación y Universidades. C.M.S. is funded by Agencia Estatal de Investigación (CNS2022-135744).

## Author contributions

D.R. conceived the study, drafted and edited the text; L.C.-R conceived the study, collected most of the experimental data, performed computational analysis and drafted the manuscript; C.M.-S. designed, collected and analysed MS data, and edited the manuscript; M.V.B.-C. collected experimental data and edited the text; D.P. collected MS data and edited the text; J.H. performed extraction and analysis of lipopeptides; M.P. and M.B.-A. designed and supervised imaging analysis and substantially edited the manuscript; J.M.-A. designed and supervised herbivore assays and edited the text; A.d.V. substantially revised and edited the text; P.C.D. substantially revised and edited the text.

## Data availability

All the raw RNA-seq data have been submitted to the Gene Expression Omnibus (GEO) and can be accessed through GEO series accession GSE299630. Metabolomics data are deposited at https://massive.ucsd.edu/ with identifier MSV000098118 All data are available within this article and its supporting information.

## References

1. Martin, F. M., Uroz, S. & Barker, D. G. Ancestral alliances: Plant mutualistic symbioses with fungi and bacteria. Science (1979) 356, (2017).

2. Soto, M. J., Domínguez-Ferreras, A., Pérez-Mendoza, D., Sanjuán, J. & Olivares, J. Mutualism *versus* pathogenesis: The give-and-take in plant-bacteria interactions. Cell Microbiol 11, 381–388 (2009).

3. Mesny, F., Hacquard, S. & Thomma, B. P. Co-evolution within the plant holobiont drives host performance. EMBO Rep 24, (2023).

4. Berg, G. & Raaijmakers, J. M. Saving seed microbiomes. ISME Journal 12, 1167–1170 (2018).

5. Shade, A., Jacques, M. A. & Barret, M. Ecological patterns of seed microbiome diversity, transmission, and assembly. Curr Opin Microbiol 37, 15–22 (2017).

6. Abdelfattah, A., Tack, A. J. M., Lobato, C., Wassermann, B. & Berg, G. From seed to seed: the role of microbial inheritance in the assembly of the plant microbiome. Trends Microbiol 31, 346–355 (2023).

7. Tsotetsi, T., Nephali, L., Malebe, M. & Tugizimana, F. Bacillus for Plant Growth Promotion and Stress Resilience: What Have We Learned? Plants 11, (2022).

8. Blake, C., Christensen, M. N. & Kovacs, A. T. Molecular aspects of plant growth promotion and protection by bacillus subtilis. Molecular Plant-Microbe Interactions 34, 15–25 (2021).

9. Berendsen, R. L. et al. Disease-induced assemblage of a plant-beneficial bacterial consortium. ISME Journal 12, 1496–1507 (2018).

10. Abd El-Daim, I. A., Bejai, S. & Meijer, J. *Bacillus velezensis* 5113 Induced Metabolic and Molecular Reprogramming during Abiotic Stress Tolerance in Wheat. Sci Rep 9, (2019).

11. Ma, K. W. et al. Coordination of microbe–host homeostasis by crosstalk with plant innate immunity. Nat Plants 7, 814–825 (2021).

12. Laurich, J. R., Lash, E., O’Brien, A. M., Pogoutse, O. & Frederickson, M. E. Community interactions among microbes give rise to host-microbiome mutualisms in an aquatic plant. mBio 15, (2024).

13. Berlanga-Clavero, M. V., et al. *Bacillus subtilis* biofilm matrix components target seed oil bodies to promote growth and anti-fungal resistance in melon. Nat Microbiol 7, 1001–1015 (2022).

14. Gu, Q. et al. Bacillomycin D produced by *Bacillus amyloliquefaciens* is involved in the antagonistic interaction with the plantpathogenic fungus Fusarium graminearum. Appl Environ Microbiol 83, (2017).

15. Roquis, D. et al. Genomic impact of stress-induced transposable element mobility in Arabidopsis. Nucleic Acids Res 49, 10431–10447 (2021).

16. Ruan, J. et al. Jasmonic acid signaling pathway in plants. Int J Mol Sci 20, (2019).

17. Howe, G. A. & Jander, G. Plant immunity to insect herbivores. Annu Rev Plant Biol 59, 41–66 (2008).

18. Di Ferdinando, M., Brunetti, C., Fini, A. & Tattini, M. Flavonoids as Antioxidants in Plants Under Abiotic Stresses. in Abiotic Stress Responses in Plants 159–179 (2012). doi:10.1007/978-1-4614-0634-1_9.

19. Evans, J. R. Improving photosynthesis. Plant Physiol 162, 1780–1793 (2013).

20. Maxwell, K. & Johnson, G. N. Chlorophyll Fluorescence-*a* Practical Guide. Journal of Experimental Botany vol. 51 (2000).

21. Foyer, C. H. & Shigeoka, S. Understanding oxidative stress and antioxidant functions to enhance photosynthesis. Plant Physiol 155, 93–100 (2011).

22. Lawlor, D. W. & Tezara, W. Causes of decreased photosynthetic rate and metabolic capacity in water-deficient leaf cells: A critical evaluation of mechanisms and integration of processes. Ann Bot 103, 561–579 (2009).

23. Graf, A., Schlereth, A., Stitt, M. & Smith, A. M. Circadian control of carbohydrate availability for growth in *Arabidopsis* plants at night. Proc Natl Acad Sci U S A 107, 9458–9463 (2010).

24. Wu, L. & Birch, R. G. Isomaltulose is actively metabolized in plant cells. Plant Physiol 157, 2094–2101 (2011).

25. Fernie, A. R., Roessner, U. & Geigenberger, P. The Sucrose Analog Palatinose Leads to a Stimulation of Sucrose Degradation and Starch Synthesis When Supplied to Discs of Growing Potato Tubers 1. Plant Physiol 125, 1967–1977 (2001).

26. Thalmann, M. et al. Regulation of leaf starch degradation by abscisic acid is important for osmotic stress tolerance in plants. Plant Cell 28, 1860–1878 (2016).

27. Kibler, C. L. et al. Evapotranspiration regulates leaf temperature and respiration in dryland vegetation. Agric For Meteorol 339, (2023).

28. Hubbart, S., et al. Enhanced thylakoid photoprotection can increase yield and canopy radiation use efficiency in rice. Commun Biol 1, (2018).

29. Sadak, M. S. & Ramadan, A. A. E. M. Impact of melatonin and tryptophan on water stress tolerance in white lupine (*Lupinus termis* L.). Physiology and Molecular Biology of Plants 27, 469–481 (2021).

30. Barón, M., Moreno-Martín, M. T. & Pineda, M. Uncovering *Botrytis cinerea*- induced physiological changes in melon plants using multi-sensor imaging approaches. Plant Stress 15, (2025).

31. Gitelson, A. A. et al. Leaf Chlorophyll Fluorescence Corrected for Re- Absorption by Means of Absorption and Reflectance Measurements*\**. Plant PhysioL WlL vol. 152 (1998).

32. Erb, M. & Kliebenstein, D. J. Plant Secondary Metabolites as Defenses, Regulators, and Primary Metabolites: The Blurred Functional Trichotomy. Plant Physiol 184, 39–52 (2020).

33. Kant, M. R., Bleeker, P. M., Wijk, M. Van, Schuurink, R. C. & Haring, M. A. Plant Volatiles in Defence. Advances in Botanical Research vol. 51 613– 666 (2009).

34. Wink, M., Schmeller, T. & Latz-Brüning, B. Modes of action of allelochemical alkaloids: Interaction with neuroreceptors, DNA, and other molecular targets. J Chem Ecol 24, 1881–1937 (1998).

35. Li, B. et al. Leaf Beetle Symbiotic Bacteria Degrade Chlorogenic Acid of Poplar Induced by Egg Deposition to Enhance Larval Survival. Plant Cell Environ (2025) doi:10.1111/pce.15427.

36. Grbić, M. et al. The genome of *Tetranychus urticae* reveals herbivorous pest adaptations. Nature 479, 487–492 (2011).

37. Alba, J. M. et al. Spider mites suppress tomato defenses downstream of jasmonate and salicylate independently of hormonal crosstalk. New Phytologist 205, 828–840 (2015).

38. Han, Z., Xiong, D., Schneiter, R. & Tian, C. The function of plant PR1 and other members of the CAP protein superfamily in plant–pathogen interactions. Mol Plant Pathol 24, 651–668 (2023).

39. Stenzel, I. et al. Jasmonate biosynthesis and the allene oxide cyclase family of *Arabidopsis thaliana*. Plant Mol Biol 51, 895–911 (2003).

40. Beyer, S. F. et al. Disclosure of salicylic acid and jasmonic acid-responsive genes provides a molecular tool for deciphering stress responses in soybean. Sci Rep 11, (2021).

41. Grifé-Ruiz, M., Hierrezuelo-León, J., de Vicente, A., Pérez-García, A. & Romero, D. Diversification of Lipopeptide Analogues Drives Versatility in Biological Activities. J Agric Food Chem 73, (2025).

42. Crisp, P. A., Ganguly, D., Eichten, S. R., Borevitz, J. O. & Pogson, B. J. Reconsidering plant memory: Intersections between stress recovery, RNA turnover, and epigenetics. Sci Adv 2, (2016).

43. Wu, W. & Fan, G. The role of epigenetics in plant pathogens interactions under the changing environments: A systematic review. Plant Stress 15, (2025).

44. Hannan Parker, A., Wilkinson, S. W. & Ton, J. Epigenetics: a catalyst of plant immunity against pathogens. New Phytologist 233, 66–83 (2022).

45. Mierziak, J. & Wojtasik, W. Epigenetic weapons of plants against fungal pathogens. BMC Plant Biol 24, (2024).

46. Mechri, B., Tekaya, M., Hammami, M. & Chehab, H. Effects of drought stress on phenolic accumulation in greenhouse-grown olive trees (Olea europaea). Biochem Syst Ecol 92, (2020).

47. Kant, M. R. et al. Mechanisms and ecological consequences of plant defence induction and suppression in herbivore communities. Ann Bot 115, 1015–1051 (2015).

48. Bolland, H. R., Gutierrez, J., Flechtmann Brill, C. H. W. & Boston·Koln, L. World Catalogue of the Spider Mite Family (Acari : Tetranychidae). (1998).

49. Njiru, C. et al. Intradiol ring cleavage dioxygenases from herbivorous spider mites as a new detoxification enzyme family in animals. BMC Biol 20, (2022).

50. Chafi, R. et al. Competitor Displacement by an Herbivore that Manipulates Plant Defences. Preprint at 10.1101/2024.05.20.594407 (2024).

51. Singh, D. P., et al. Roots of resistance: Unraveling microbiome-driven plant immunity. Plant Stress 14, (2024).

52. Pirttilä, A. M. et al. Exchange of Microbiomes in Plant-Insect Herbivore Interactions. mBio 14, (2023).

53. Wang, X., Li, Y., Rensing, C. & Zhang, X. Early inoculation and bacterial community assembly in plants: A review. Microbiol Res 296, (2025).

54. Hadian, S., Smith, D. L. & Supronienė, S. Modulating the Plant Microbiome: Effects of Seed Inoculation with Endophytic Bacteria on Microbial Diversity and Growth Enhancement in Pea Plants. Microorganisms 13, (2025).

55. Kant, M. R., Sabelis, M. W., Haring, M. A. & Schuurink, R. C. Intraspecific variation in a generalist herbivore accounts for differential induction and impact of host plant defences. Proceedings of the Royal Society B: Biological Sciences 275, 443–452 (2008).

56. Ontiveros, I., López-Moya, J. J. & Díaz-Pendón, J. A. Coinfection of Tomato Plants with Tomato yellow leaf curl virus and Tomato chlorosis virus Affects the Interaction with Host and Whiteflies. Phytopathology 112, 741–975 (2022).

57. Schneider, C. A., Rasband, W. S. & Eliceiri, K. W. NIH Image to ImageJ: 25 years of image analysis. Nat Methods 9, 671–675 (2012).

58. Ruggieri, V. et al. An improved assembly and annotation of the melon (Cucumis melo L.) reference genome. Sci Rep 8, (2018).

59. Sysi-Aho, M., Katajamaa, M., Yetukuri, L. & Orešič, M. Normalization method for metabolomics data using optimal selection of multiple internal standards. BMC Bioinformatics 8, (2007).

60. Petras, D. et al. Mass spectrometry-based visualization of molecules associated with human habitats. Anal Chem 88, 10775–10784 (2016).

61. Schmid, R. et al. Integrative analysis of multimodal mass spectrometry data in MZmine 3. Nat Biotechnol 41, 447–449 (2023).

62. Wang, M. et al. Sharing and community curation of mass spectrometry data with Global Natural Products Social Molecular Networking. Nat Biotechnol 34, 828–837 (2016).

63. Shannon, P. et al. Cytoscape: A software Environment for integrated models of biomolecular interaction networks. Genome Res 13, 2498–2504 (2003).

64. Dührkop, K. et al. SIRIUS 4: a rapid tool for turning tandem mass spectra into metabolite structure information. Nat Methods 16, 299–302 (2019).

65. Kim, H. W. et al. NPClassifier: A Deep Neural Network-Based Structural Classification Tool for Natural Products. J Nat Prod 84, 2795–2807 (2021).

66. Djoumbou Feunang, Y., et al. ClassyFire: automated chemical classification with a comprehensive, computable taxonomy. J Cheminform 8, 1–20 (2016).

67. Sumner, L. W. et al. Proposed minimum reporting standards for chemical analysis: Chemical Analysis Working Group (CAWG) Metabolomics Standards Initiative (MSI). Metabolomics 3, 211–221 (2007).

68. Pang, Z. et al. MetaboAnalystR 4.0: a unified LC-MS workflow for global metabolomics. Nat Commun 15, (2024).

69. Protsyuk, I. et al. 3D molecular cartography using LC-MS facilitated by Optimus and ’ili software. Nat Protoc 13, 134–154 (2018).

70. Pineda, M., Soukupová, J., Matouš, K., Nedbal, L. & Barón, M. Conventional and combinatorial chlorophyll fluorescence imaging of tobamovirus-infected plants. Photosynthetica 46, 441–451 (2008).

71. Polonio, Á. et al. RNA-seq analysis and fluorescence imaging of melon powdery mildew disease reveal an orchestrated reprogramming of host physiology. Sci Rep 9, (2019).

72. Lichtenthaler, H. K. & Buschmann, C. Chlorophylls and Carotenoids: Measurement and Characterization by UV-VIS Spectroscopy. Current Protocols in Food Analytical Chemistry 1, 1–8 (2001).

73. Pineda, M., Pérez-Bueno, M. L. & Barón, M. Novel Vegetation Indices to Identify Broccoli Plants Infected With *Xanthomonas campestris* pv. campestris. Front Plant Sci 13, (2022).

74. Pérez-Bueno, M. L., Pineda, M., Díaz-Casado, E. & Barón, M. Spatial and temporal dynamics of primary and secondary metabolism in *Phaseolus vulgaris* challenged by *Pseudomonas syringae*. Physiol Plant 153, 161– 174 (2015).

75. Ontiveros, I., López-Moya, J. J. & Díaz-Pendón, J. A. Coinfection of Tomato Plants with Tomato yellow leaf curl virus and Tomato chlorosis virus Affects the Interaction with Host and Whiteflies. Phytopathology 112, 944–952 (2022).

76. Tamilselvan, R., Mohan Kumar, S., Mahalingam, C. & Senguttuvan, K. Biology of whitefly, *Bemisia tabaci* (Genn.) (Hemiptera, Aleyrodidae) on resistant and susceptible cotton genotypes. J Entomol Zool Stud 7, 644– 647 (2019).

77. Michán, C. & Pueyo, C. Growth phase-dependent variations in transcript profiles for thioredoxin- and glutathione-dependent redox systems followed by budding and hyphal *Candida albicans* cultures. FEMS Yeast Res 9, 1078–1090 (2009).

78. Livak, K. J. & Schmittgen, T. D. Analysis of relative gene expression data using real-time quantitative PCR and the 2-ΔΔCT method. Methods 25, 402–408 (2001).

