## Supplementary Figures and Tables for "Early seed priming with closely related *Bacillus* strains induces divergent physiological and defense responses in melon"

#### Figure S1

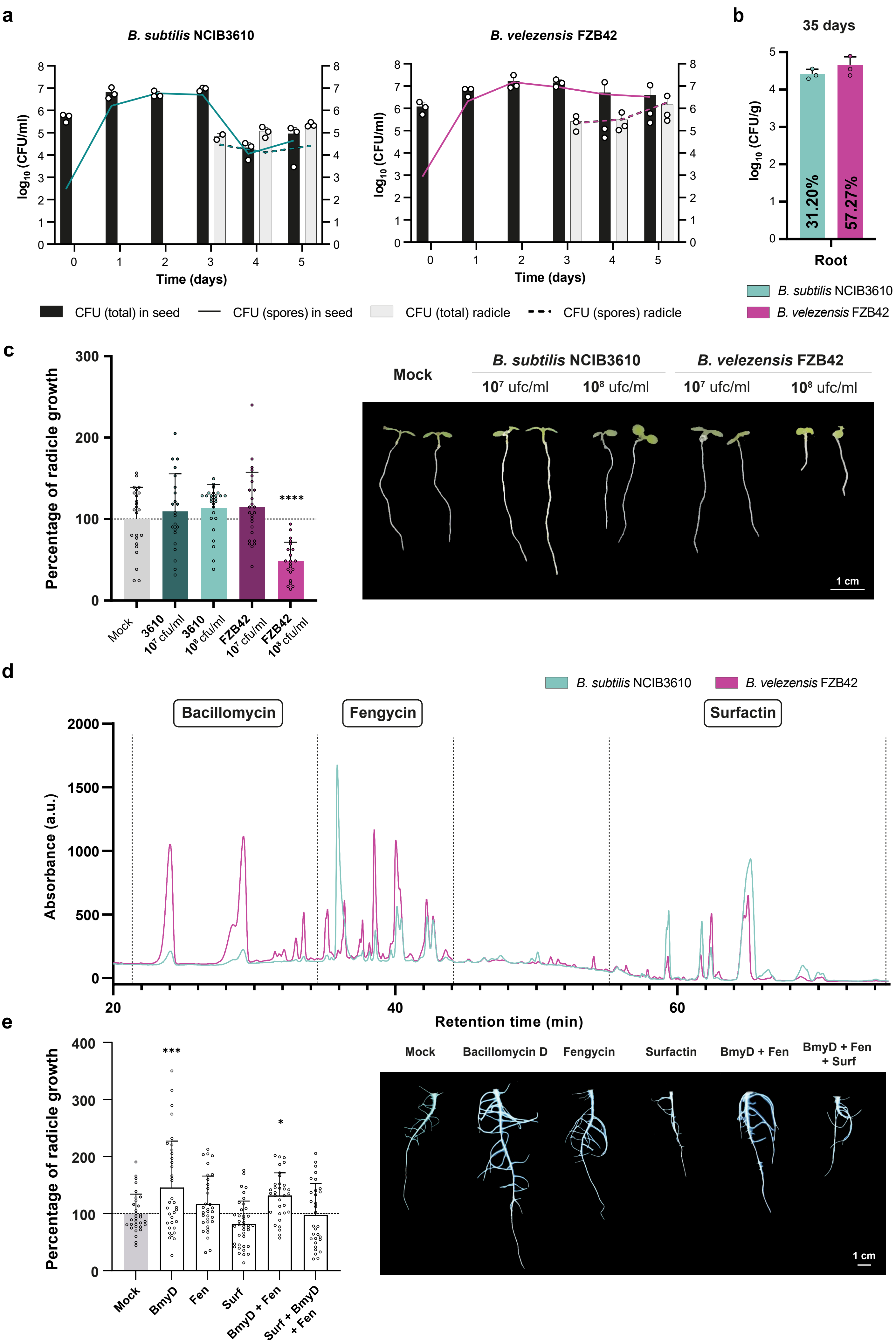

a
B. subtilis NCIB3610
B. velezensis FZB42
log10 (CFU/ml)
Time (days)
CFU (total) in seed
CFU (spores) in seed
CFU (total) radicle
CFU (spores) radicle
b
35 days
log10 (CFU/g)
31.20%
57.27%
Root
B. subtilis NCIB3610
B. velezensis FZB42
c
Percentage of radicle growth
****
Mock
3610 10^7 cfu/ml
3610 10^8 cfu/ml
FZB42 10^7 cfu/ml
FZB42 10^8 cfu/ml
Mock
B. subtilis NCIB3610
B. velezensis FZB42
10^7 ufc/ml
10^8 ufc/ml
10^7 ufc/ml
10^8 ufc/ml
1 cm
d
B. subtilis NCIB3610
B. velezensis FZB42
Bacillomycin
Fengycin
Surfactin
Absorbance (a.u.)
Retention time (min)
e
Percentage of radicle growth
***
*
Mock
BmyD
Fen
Surf
BmyD + Fen
Surf + BmyD + Fen
Mock
Bacillomycin D
Fengycin
Surfactin
BmyD + Fen
BmyD + Fen + Surf
1 cm

**Figure S1. Dose-dependent effects of *Bacillus* on early development and metabolite production.** **a** Dynamics of CFU ($\log_{10}$ CFU/ml) and sporulation percentage of *B. subtilis* (left) and *B. velezensis* (right) in seed and radicle extracts over 5 days post-treatment. **b** Total CFU/g in root tissue and average sporulation rate in plants grown from treated seeds. **c** Left: Radicle growth (%) 5 days after treatment of *Arabidopsis thaliana* seeds with each strain at two inoculum densities. Data normalized to mock controls. Mean ± SD. One-way ANOVA with Dunnett's test (***P < 0.0001). Right: Representative radicle images. Scale bar = 1 cm. **d** HPLC chromatograms of major cyclic lipopeptides in SPE-purified extracts from both strains. **e** Left: Radicle growth (%) after treatment with bacillomycin D, fengycin, and surfactin (20 μM), applied individually or in combination. Mean ± SD. Dunnett's test (*P = 0.0199; ***P = 0.0007). Right: Representative radicles. Scale bar = 1 cm.

**Figure S2**

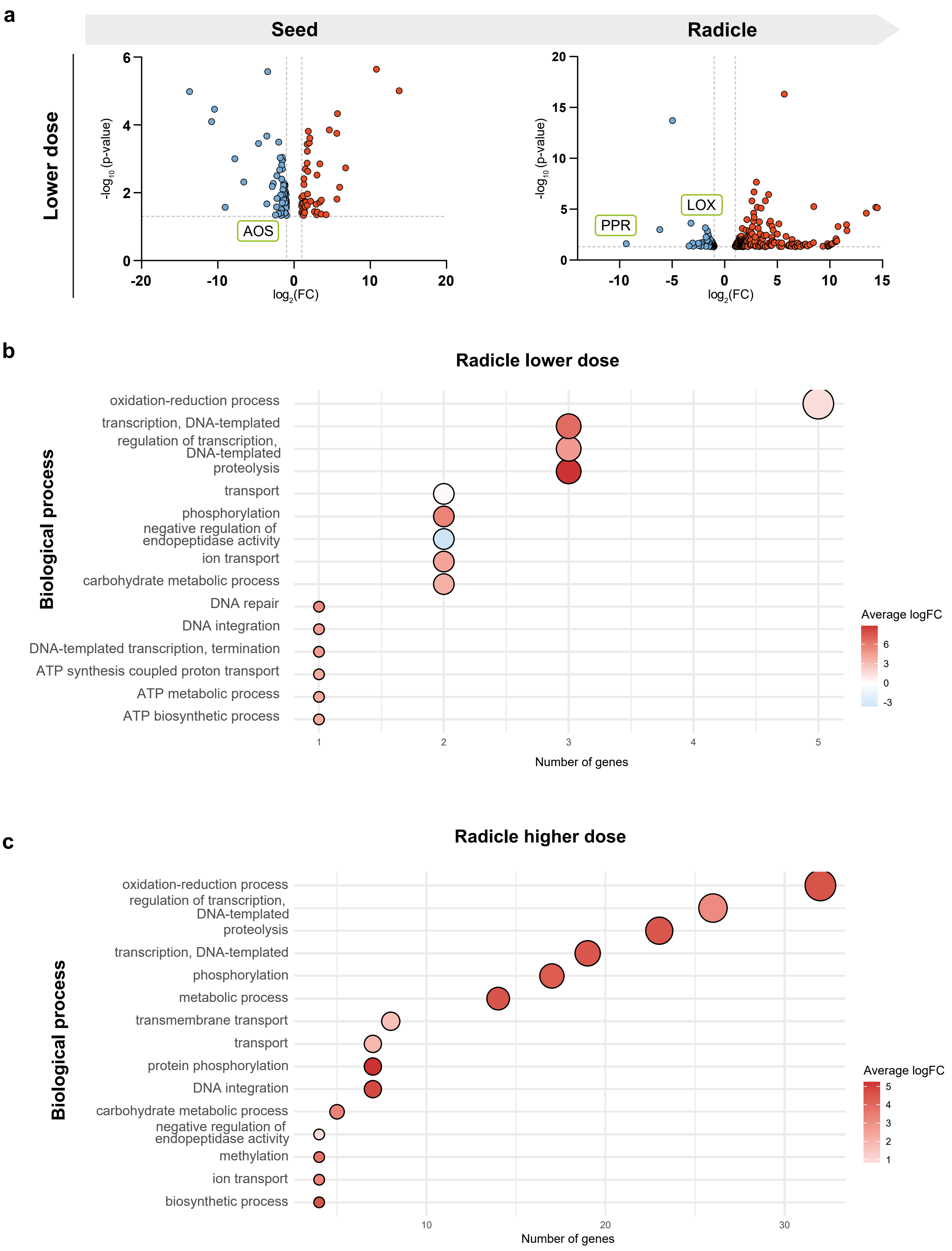

a
Seed
Radicle
Lower dose
-log10 (p-value)
log2(FC)
AOS
LOX
PPR
b
Radicle lower dose
Biological process
oxidation-reduction process
transcription, DNA-templated
regulation of transcription, DNA-templated
proteolysis
transport
phosphorylation
negative regulation of endopeptidase activity
ion transport
carbohydrate metabolic process
DNA repair
DNA integration
DNA-templated transcription, termination
ATP synthesis coupled proton transport
ATP metabolic process
ATP biosynthetic process
Number of genes
Average logFC
c
Radicle higher dose
Biological process
oxidation-reduction process
regulation of transcription, DNA-templated
proteolysis
transcription, DNA-templated
phosphorylation
metabolic process
transmembrane transport
transport
protein phosphorylation
DNA integration
carbohydrate metabolic process
negative regulation of endopeptidase activity
methylation
ion transport
biosynthetic process
Number of genes
Average logFC

**Figure S2. Low-dose *B. velezensis* treatment reduces transcriptional and oxidative activity. a** Volcano plot of DEGs in seeds and radicles treated with low-dose *B. velezensis*. Thresholds shown as dashed lines. Tags highlight genes related to retrotransposon activity, oxidative stress, and defense. **b–c** GO enrichment (Biological Process) analysis for radicle DEGs after low (b) and high (c) *B. velezensis* treatment.

**Figure S3**

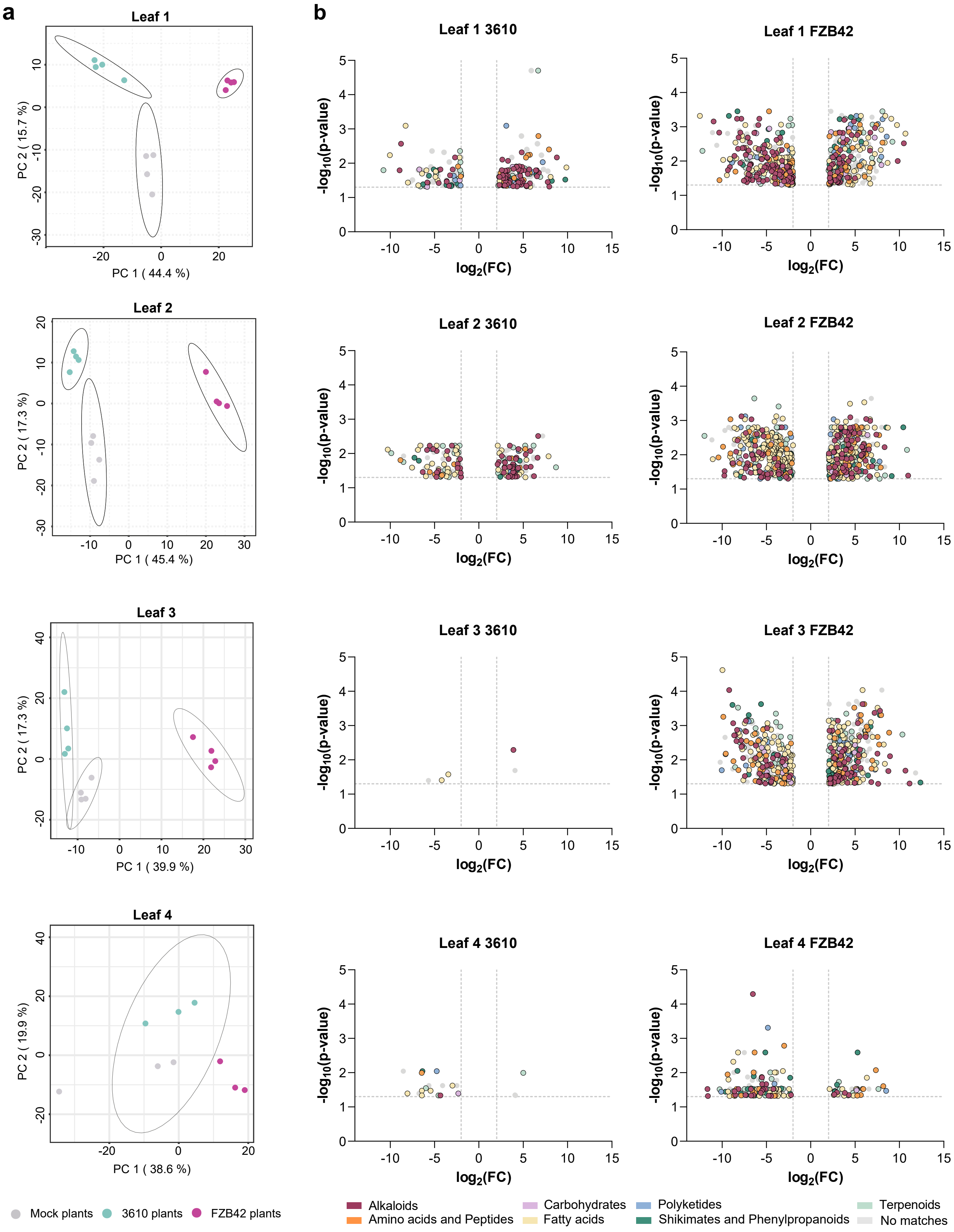
a
Leaf 1
PC 1 ( 44.4 %)
PC 2 ( 15.7 %)
Leaf 2
PC 1 ( 45.4 %)
PC 2 ( 17.3 %)
Leaf 3
PC 1 ( 39.9 %)
PC 2 ( 17.3 %)
Leaf 4
PC 1 ( 38.6 %)
PC 2 ( 19.9 %)
Mock plants
3610 plants
FZB42 plants
b
Leaf 1 3610
Leaf 1 FZB42
Leaf 2 3610
Leaf 2 FZB42
Leaf 3 3610
Leaf 3 FZB42
Leaf 4 3610
Leaf 4 FZB42
-log10(p-value)
log2(FC)
Alkaloids
Amino acids and Peptides
Carbohydrates
Fatty acids
Polyketides
Shikimates and Phenylpropanoids
Terpenoids
No matches

**Figure S3.** ***B. velezensis*** **induces the strongest shifts in leaf metabolome across developmental stages. a** PCA of metabolomic profiles across leaf age (leaf 1 = oldest to leaf 4 = youngest). **b** Volcano plots showing differentially accumulated metabolites ($log_2FC > 2$, $FDR < 0.05$) by treatment and leaf age. Colors indicate chemical classes (NPC#pathway).

#### Figure S4

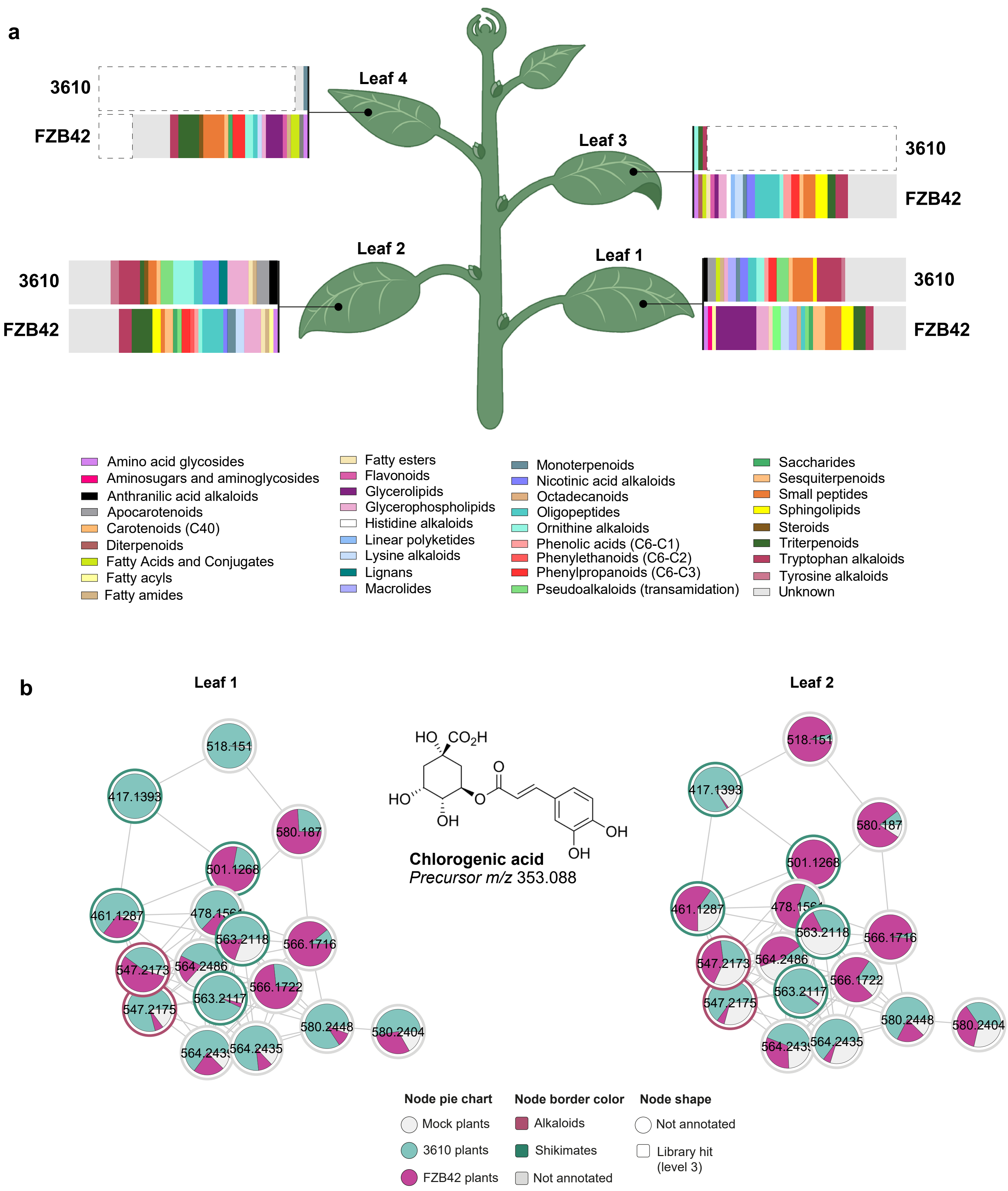
a
3610
FZB42
Leaf 4
Leaf 3
Leaf 2
Leaf 1
Amino acid glycosides
Aminosugars and aminoglycosides
Anthranilic acid alkaloids
Apocarotenoids
Carotenoids (C40)
Diterpenoids
Fatty Acids and Conjugates
Fatty acyls
Fatty amides
Fatty esters
Flavonoids
Glycerolipids
Glycerophospholipids
Histidine alkaloids
Linear polyketides
Lysine alkaloids
Lignans
Macrolides
Monoterpenoids
Nicotinic acid alkaloids
Octadecanoids
Oligopeptides
Ornithine alkaloids
Phenolic acids (C6-C1)
Phenylethanoids (C6-C2)
Phenylpropanoids (C6-C3)
Pseudoalkaloids (transamidation)
Saccharides
Sesquiterpenoids
Small peptides
Sphingolipids
Steroids
Triterpenoids
Tryptophan alkaloids
Tyrosine alkaloids
Unknown
b
Leaf 1
Leaf 2
Chlorogenic acid
Precursor m/z 353.088
Node pie chart
Mock plants
3610 plants
FZB42 plants
Node border color
Alkaloids
Shikimates
Not annotated
Node shape
Not annotated
Library hit (level 3)

**Figure S4. Specific classes of metabolites accumulate in *Bacillus*-primed plants. a** Classification of the 50 most significantly accumulated metabolites ($log_2FC > 2$, FDR < 0.05) by leaf and treatment, grouped by NPC#superclass. **b** Molecular family network of a chlorogenic acid analog differentially abundant in the first two leaves. Pie charts show mean relative abundance. Node shape: identification confidence (Sumner et al., 2007); border: NPC#pathway class.

#### Figure S5

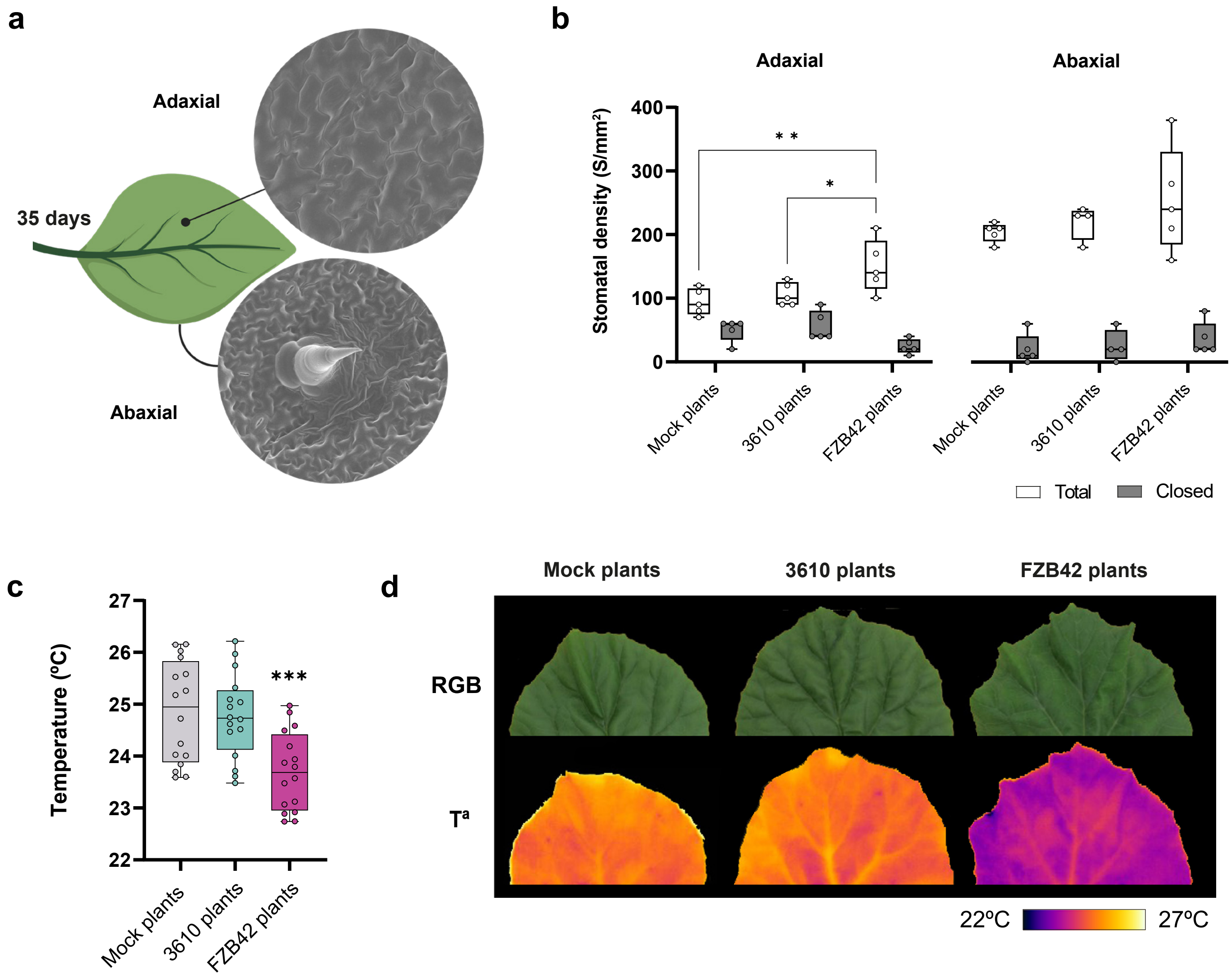

a
Adaxial
35 days
Abaxial
b
Adaxial
Abaxial
Stomatal density (S/mm²)
400
300
200
100
0
* *
*
Mock plants
3610 plants
FZB42 plants
Mock plants
3610 plants
FZB42 plants
Total
Closed
c
Temperature (ºC)
27
26
25
24
23
22
***
Mock plants
3610 plants
FZB42 plants
d
Mock plants
3610 plants
FZB42 plants
RGB
Tª
22ºC
27ºC

**Figure S5.** ***B. velezensis*****-treated plants show increased stomatal density and altered leaf temperature. a** SEM images of adaxial (top) and abaxial (bottom) leaf surfaces. **b** Stomatal density (stomata/mm$^2$) by surface and treatment. Two-way ANOVA with Tukey's test (*P = 0.0210; **P = 0.0032). **c** Surface temperature (°C) of adaxial leaf surface. One-way ANOVA with Dunnett's test (***P = 0.0009). **d** Representative RGB and thermal images of adaxial surfaces.

### Figure S6

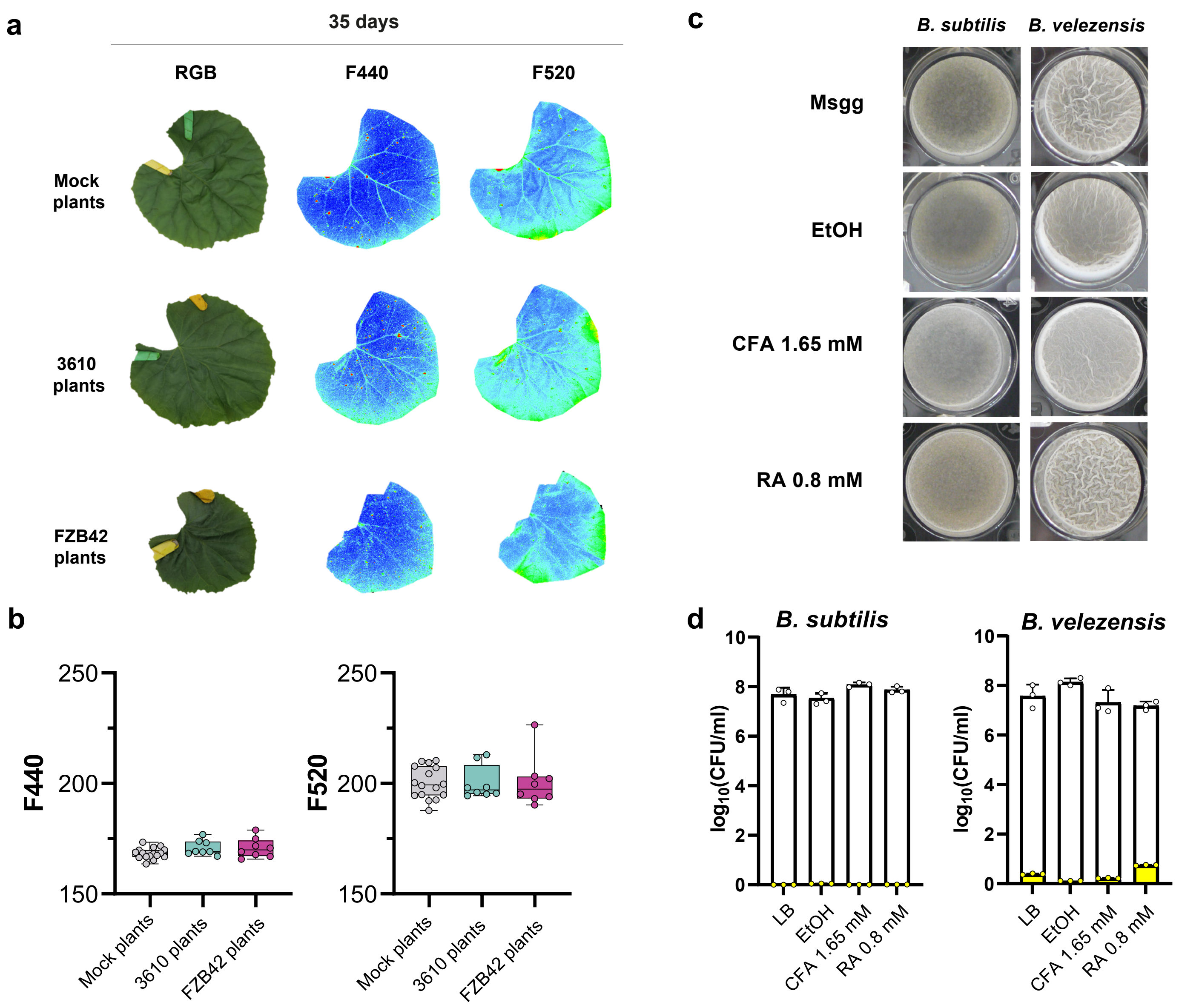
a
35 days
RGB
F440
F520
Mock plants
3610 plants
FZB42 plants
b
F440
F520
250
200
150
Mock plants
3610 plants
FZB42 plants
c
B. subtilis
B. velezensis
Msgg
EtOH
CFA 1.65 mM
RA 0.8 mM
d
B. subtilis
B. velezensis
log10(CFU/ml)
LB
EtOH
CFA 1.65 mM
RA 0.8 mM
CFU Vegetative
CFU Spores

**Figure S6. Leaf-soluble phenolics are non-cytotoxic to *Bacillus* and absent in healthy tissues. a** Fluorescence images (F440, F520) of phenolic compounds in cell walls. **b** Quantified fluorescence emission from leaves by treatment. **c** Biofilm phenotypes of both *Bacillus* strains in Msgg media with caffeic acid (CFA) or rosmarinic acid (RA); ethanol control included. **d** CFU/ml of strains after 24 h exposure to CFA and RA in LB; control = 0.8% ethanol.

**Figure S7**

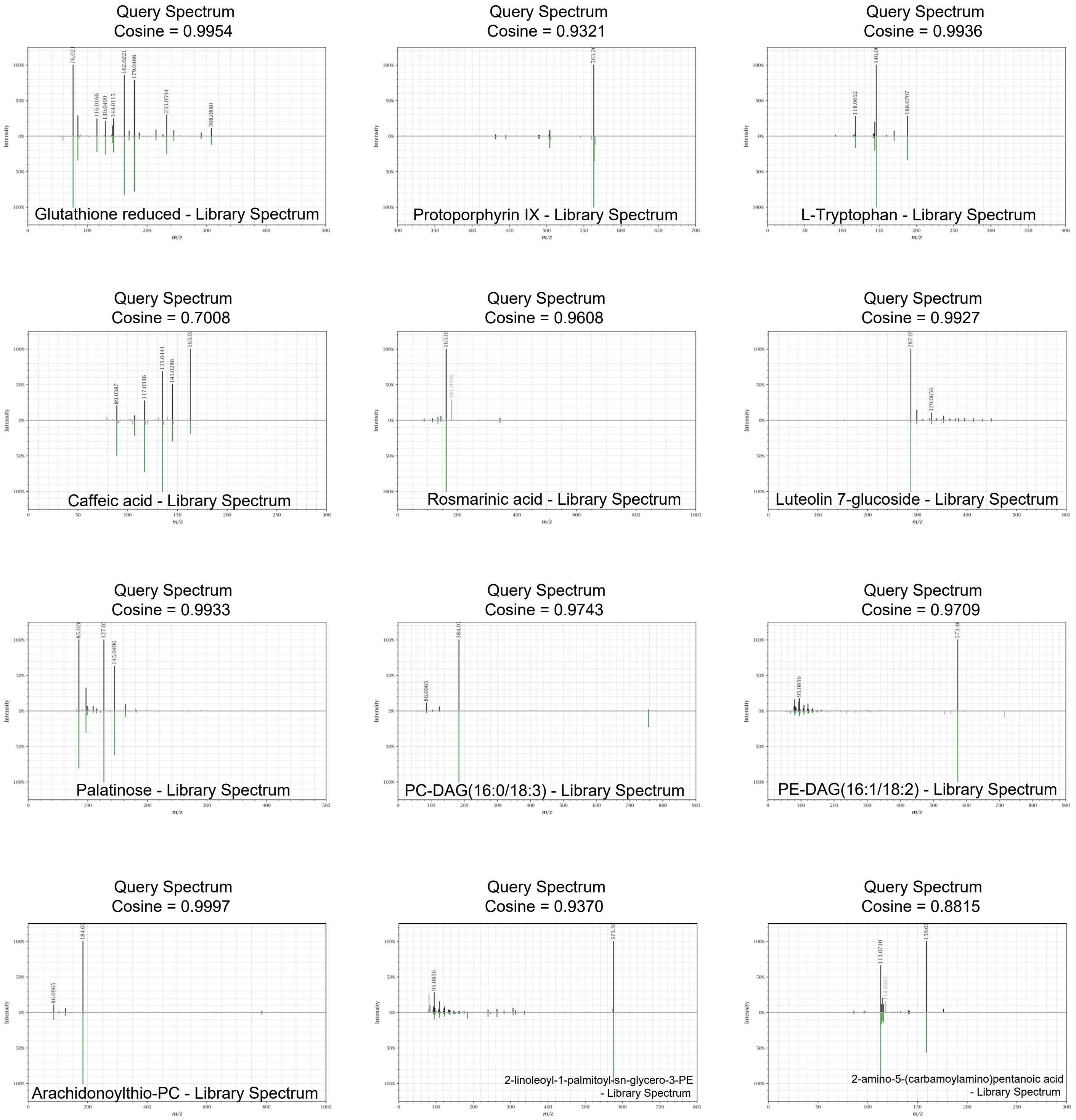
Query Spectrum
Cosine = 0.9954
Glutathione reduced - Library Spectrum
Query Spectrum
Cosine = 0.9321
Protoporphyrin IX - Library Spectrum
Query Spectrum
Cosine = 0.9936
L-Tryptophan - Library Spectrum
Query Spectrum
Cosine = 0.7008
Caffeic acid - Library Spectrum
Query Spectrum
Cosine = 0.9608
Rosmarinic acid - Library Spectrum
Query Spectrum
Cosine = 0.9927
Luteolin 7-glucoside - Library Spectrum
Query Spectrum
Cosine = 0.9933
Palatinose - Library Spectrum
Query Spectrum
Cosine = 0.9743
PC-DAG(16:0/18:3) - Library Spectrum
Query Spectrum
Cosine = 0.9709
PE-DAG(16:1/18:2) - Library Spectrum
Query Spectrum
Cosine = 0.9997
Arachidonoylthio-PC - Library Spectrum
Query Spectrum
Cosine = 0.9370
2-linoleoyl-1-palmitoyl-sn-glycero-3-PE
- Library Spectrum
Query Spectrum
Cosine = 0.8815
2-amino-5-(carbamoylamino)pentanoic acid
- Library Spectrum

**Figure S7. Mirror plot comparison of candidate MS/MS spectra to GNPS reference standards.** Upper trace: candidate metabolite (black); lower trace: standard compound (green). Generated via https://metabolomics-usi.ucsd.edu.

**Table S1. Differential expression ($log_2$ fold change) of selected genes in seeds and radicles treated with low and high doses of *B. velezensis*. Ordered by gene ID.**

| | | | Seed | | Radicle | |
|---|---|---|---|---|---|---|
| ID | Description | Abbr. | Low | High | Low | High |
| MELO3C002457 | Peroxidase | Perox. | - | -1.605 | - | -1.048 |
| MELO3C005540 | 14 kDa proline-rich protein DC2.15-like | PRP | - | -1.488 | - | - |
| MELO3C008512 | Pentatricopeptide repeat-containing protein | PPR | - | - | -9.367 | 1.917 |
| MELO3C009108 | Non-specific lipid-transfer protein | NSLTP | -1.168 | -1.593 | -1.218 | - |
| MELO3C011383 | DNA-directed RNA polymerase subunit beta | RNApol subB | -4.651 | -3.119 | 2.918 | - |
| MELO3C014217 | HTH myb-type domain-containing protein | HTH | - | 2.26 | 1.318 | - |
| MELO3C014630 | Lipoxygenase | LOX | - | - | -2.981 | 2.46 |
| MELO3C018413 | Allene oxide synthase | AOS | -1.717 | -2.891 | - | - |
| MELO3C020689 | Respiratory burst oxidase | RBOH | - | - | - | 7.002 |
| MELO3C027069 | Proteasome subunit beta | Prot. | - | -13.095 | 14.356 | 13.849 |
| MELO3C027194 | DNA-directed RNA polymerase II subunit RPB7 | RNApol subR | - | -10.823 | 10.766 | 10.222 |
| MELO3C027784 | Gag/pol protein | Gag/pol | - | 9.693 | - | - |
| MELO3C028181 | Gag/pol protein | Gag/pol | - | 9.242 | - | - |
| MELO3C028271 | Plant transposase | Plant trans. | - | - | - | 8.649 |
| MELO3C031213 | Gag/pol protein | Gag/pol | - | - | - | 8.79 |
| MELO3C034502 | Gag/pol polyprotein | Gag/pol | - | 8.544 | - | - |

**Table S2**. **Representative metabolites from *B. subtilis*-treated plants with identification parameters per Sumner et *al.*, 2007**.

| Proposed Compound Name | Leaf | $log_2$ FC | IonMode | Adduct | Observed m/z | Calculated m/z | Ppm error |
|---|---|---|---|---|---|---|---|
| Rosmarinic acid | 1 | 8.2188 | Positive | $[M+H]^+$ | 343.1024 | 343.083 | 56.4839 |
| Caffeic acid | 1 / 2 | 4.6769 / 3.0102 | Positive | $[M+H]^+$ | 163.0391 | 181.056 | 0.8423 |
| Palatinose | 1 | 3.4228 | Positive | $[M+Na]^+$ | 325.0917 | 325.111 | 59.3247 |
| Abrine | 1 / 2 | 4.4624 / 3.5593 | Positive | $[M+H]^+$ | 188.0709 | 188.07 | 4.7869 |
| L-Tryptophan | 1 / 2 | 6.5941 / 7.1977 | Positive | M+H | 205.0976 | 205.097 | 2.9015 |
| Protoporphyrin IX | 1 / 2 / 3 | -4.3462 / -5.548 / -3.5456 | Positive | $[M+H]^+$ | 563.2659 | 563.253 | 22.9727 |
| PC-DAG (16:0/18:3) | 2 | 4.5576 | Positive | $[M+H]^+$ | 756.5539 | 756.551 | 3.7917 |
| Isoorientin | 2 | 1.7423 | Positive | $[M+H]^+$ | 449.1083 | 449.108 | 1.0193 |

**Table S3**. **Representative metabolites from *B. velezensis*-treated plants with identification parameters per Sumner et *al*., 2007**.

| Metabolite | Leaf | $log_2$ FC | IonMode | Adduct | Observed m/z | Calculated m/z | Ppm error |
|---|---|---|---|---|---|---|---|
| 2-Linoleoyl-1-palmitoyl-sn-glycero-3-phosphoetanolamine | 1 / 2 | 7.8944 / 5.987 | Positive | $[M+H]^+$ | 716.5226 | 716.523 | 0.59628 |
| 2-amino-5-(carbamoylamino)pentanoic acid (citrulline) | 1 | 6.6909 | Positive | $[M+H]^+$ | 176.1037 | 176.103 | 3.9858 |
| Isoorientin | 1 / 2 | 5.4093 / 2.6828 | Positive | $[M+H]^+$ | 449.1083 | 449.108 | 1.0193 |
| Isovitexin | 1 | 4.524 | Positive | $[M+H]^+$ | 433.1132 | 433.113 | 0.5637 |
| Arachidonoylthio-PC | 1 | 4.4946 | Positive | M+H | 784.5885 | 784.584 | 5.7567 |
| PC-DAG (16:0/18:3) | 1 / 2 | 4.4863 / 5.3095 | Positive | $[M+H]^+$ | 756.5539 | 756.551 | 3.7917 |
| PE-DAG (16:1/18:2) | 1 / 2 | 3.8627 / 3.8916 | Positive | $[M+H]^+$ | 714.5075 | 714.506 | 2.1356 |
| Saponarin | 1 | 3.281 | Positive | $[M+H]^+$ | 595.166 | 595.166 | 0.4102 |
| Luteolin-7-glucoside | 1 | 3.064 | Positive | $[M+H]^+$ | 449.1084 | 449.108 | 0.8834 |
| Glutathione reduced | 1 | 2.7304 | Positive | $[M+H]^+$ | 308.0909 | 308.091 | 0.2972 |
| Protoporphyrin IX | 2 / 3 | -3.6758 / -2.8864 | Positive | $[M+H]^+$ | 563.2659 | 563.253 | 22.9727 |
| 1,2-Ditetradecanoyl-sn-glycero-3-phosphocholine | 2 / 3 | 8.3261 / 8.7569 | Positive | $[M+H]^+$ | 678.4735 | 678.474 | 0.7197 |

| | | | | | | | |
|---|---|---|---|---|---|---|---|
| L-Tryptophan | 2 / 3 | 6.0517 / 6.18 | Positive | M+H | 205.0976 | 205.097 | 2.9015 |
| 3-Indoleacrylic acid | 2 / 3 | 4.3084 / 4.3484 | Positive | M+H-$H_2O$ | 170.0604 | 170.06 | 2.1534 |
| Sinapic acid | 2 / 3 / 4 | 2.207 / 3.2603 / 1.2376 | Positive | $[M+H]^+$ | 207.0654 | 207.065 | 1.1054 |
| Ferulate | 3 | 3.3913 | Positive | $[M+H]^+$ | 195.0655 | 195.065 | 2.5814 |
| Abrine | 3 | 2.8197 | Positive | $[M+H]^+$ | 188.0709 | 188.07 | 4.7869 |
| Methyl trans-cinnamic acid | 3 | 2.2938 | Positive | $[M+H]^+$ | 163.0755 | 163.075 | 3.0878 |
| α-tocopherol | 3 | 1.7172 | Positive | $[M+H]^+$ | 430.3798 | 430.38 | 0.4964 |
| DL-Phenylalanine | 3 | 1.5699 | Positive | $[M+H]^+$ | 166.0865 | 166.083 | 21.1311 |

**Table S4. Primer sequences used in qRT-PCR analyses.**

| Gene | GeneID | Primers pairs | Sequence (5' → 3') | Amplicon (bp) |
|---|---|---|---|---|
| Pathogenesis-related protein PRB1-2 (PR1-1a) | 103495329 | CmPR1_F | GCATCAACGACTGTAGGCTAGT | 117 |
| | | CmPR1_R | ACTGCTTCTCATTCACCCACAT | |
| Allene Oxide Synthase (AOS) | 103487935 | Cm_AOS_fwd | CGCTACGAGGCCATCTACAG | 92 |
| | | Cm_AOS_rev | CTTCAAAGATGCCACTGCCG | |
| Allene Oxide Cyclase (AOC) | 103503379 | Cm_AOC_fwd | GGTTCAGAACAAGCAGTGCG | 121 |
| | | Cm_AOC_rev | CGAAGGAGTCGTAACGGAGG | |
| Actin 7 *(act7)* | 103485254 | CmActin7_F | CACTGGTATTGTGCTGGATTC | 98 |
| | | CmActin7_R | CAAGGTCCAAACGGAGAATG | |
